# Bioinformatic analysis of sulfotransferases from an unexplored gut microbe, *Sutterella wadsworthensis 3_1_45B*: Possible roles towards detoxification via sulfation by the members of the human gut microbiome

**DOI:** 10.1101/2024.01.08.574607

**Authors:** Lauryn Langford, Dhara D. Shah

**Author notes:** Corresponding author, (DDS).

## Abstract

Sulfation, primarily facilitated by sulfotransferases, plays a crucial role in the detoxification pathways of both endogenous substances and xenobiotics, enhancing their water solubility and promoting metabolism and elimination. Traditionally, this bioconversion has been attributed to a family of human cytosolic sulfotransferases (hSULTs) known for their high sequence similarity and dependence on 3’-phosphoadenosine 5’-phosphosulfate (PAPS) as a sulfate donor. However, recent studies have revealed the presence of PAPS-dependent sulfotransferases within gut commensals, indicating that the gut microbiome may harbor a diverse array of sulfotransferase enzymes and may contribute to detoxification processes via sulfation. In this study, we investigated the prevalence of sulfotransferases in the members of the human gut microbiome. Interestingly, we stumbled upon a different class of sulfotransferases, known as aryl-sulfate sulfotransferases (ASSTs). ASSTs have been characterized from a few different prokaryotes including *E. coli*. ASSTs do not utilize PAPS which is the default sulfate donor for the human sulfotransferases. Our bioinformatics analyses revealed that the gut microbial genus *Sutterella* possesses a significant number of *asst* genes, possibly encoding multiple ASST enzymes. Fluctuations in the microbes of the genus *Sutterella* have been associated with various health conditions. For this reason, we characterized 17 different ASSTs from *Sutterella wadsworthensis 3_1_45B* with bioinformatics. Our findings reveal that *Sw*ASSTs share similarities with *E. coli* ASST but also exhibit significant structural variations and sequence diversity. These differences might drive potential functional diversification and likely reflect an evolutionary divergence from their PAPS-dependent counterparts.

## Introduction

Sulfation is a key detoxification pathway, where endogenous compounds and foreign substances undergo a transformation to increase their water solubility, aiding in metabolism and elimination(1). This process has traditionally been attributed to a diverse array of human cytosolic sulfotransferases(2–20). However, recent discoveries have identified enzymes capable of cholesterol sulfation within the *Bacteroides* genus, a prominent member of the human gut microbiota(21, 22). This suggests the potential presence of additional sulfotransferases in the gut microbiome. While microbial sulfotransferases have been previously studied, they lack the extensive characterization seen with human enzymes(23–26). Sulfotransferases generally facilitate a biochemical reaction in which a sulfate group is transferred from a donor to an acceptor molecule (Fig 1).

**Fig 1.**
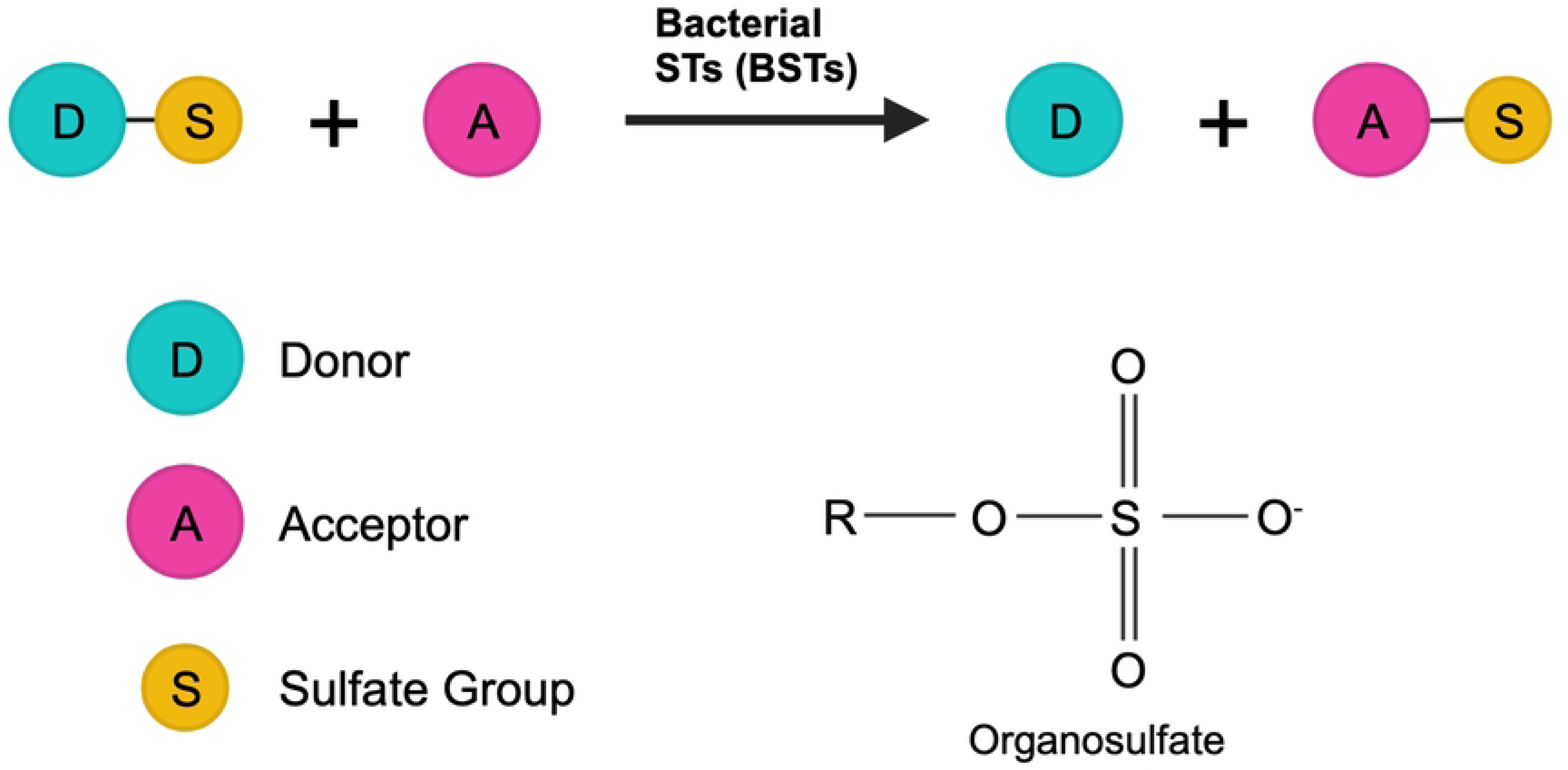
Depiction of a sulfotransferase catalyzed reaction. This scheme illustrates the general chemical reaction catalyzed by sulfotransferases where sulfate group is transferred from a donor molecule to an acceptor molecule.

Extensive research has been conducted on human sulfotransferases. To date, fourteen cytosolic human sulfotransferases (hSULTs) have been identified and characterized (S1 Table)(2–20). These enzymes exhibit a high degree of sequence similarity, and all utilize 3’-phosphoadenosine 5’-phosphosulfate (PAPS) as the common sulfate donor. However, their sulfate acceptor preferences differ, giving each hSULT a unique function(27). This variability is mostly present in the substrate binding regions of the enzymes(28). Specifically, the substrate binding loops of hSULTs demonstrate significant diversity(27). The fourteen distinct isoforms of human sulfotransferases (hSULTs) have tissue specific expression and are responsible for the sulfation of numerous small molecules, both endogenous and exogenous(28). These enzymes are typically characterized by their dependency on 3’-phosphoadenosine 5’-phosphosulfate (PAPS) for the sulfotransferase activity and are recognized by the presence of PAPS binding motifs (29). In addition, hSULTs are categorized under the large class of aryl sulfotransferases (EC 2.8.2.1)(26). Human cytosolic sulfotransferases facilitate sulfate transfer reactions, utilizing either a non-sequential (random Bi Bi) mechanism, where the substrates bind independently without intermediate formation, or a sequential (ordered Bi Bi) mechanism, where the binding of one substrate facilitates the attachment of the other (20, 30, 31). PAPS-dependent aryl sulfotransferases are not exclusive to humans but are also found across various eukaryotic and prokaryotic species(7, 9, 11, 17, 21, 22). The widespread and prevalent nature of these enzymes is not fully understood. One hypothesis is that these enzymes might be involved in detoxification of molecules that are present at varying levels in the environments of these diverse life forms(30).

In the late 1980s, a series of studies demonstrated that the intestinal flora exhibit sulfation activity on small phenolic compounds (23, 24, 26). These investigations revealed a new class of previously unknown microbial sulfotransferases, distinctive in its utilization of sulfate donors other than the mammalian default, 3’-phosphoadenosine 5’-phosphosulfate (PAPS). Interestingly, these microbial enzymes were capable of utilizing a variety of non-physiological phenolic sulfate esters as donors, with p-nitrophenol sulfate (pNPS) being predominantly used.

Due to the differences in the donor specificities, reaction mechanisms, and kinetic profiles, these enzymes were designated as aryl-sulfate sulfotransferases (ASSTs), classified under EC 2.8.2.22 (32). Beak et al. conducted studies on the distribution of ASSTs, showing their presence across both Gram-positive and Gram-negative bacterial species(33). Furthermore, ASSTs were observed to exist both with and without signal peptide (33). Cumulatively, these studies show that there is a great deal of variations in microbial ASSTs. The widespread prevalence of ASSTs and the current gap in understanding their functional importance demonstrate the need for further study.

The integral role of gut microbiota in metabolizing a range of xenobiotics and endogenous substances is already well-recognized(34). As mentioned earlier, a common route for the metabolism of these compounds is via sulfation. Motivated by these findings, we have delved into the genomic landscape of gut microbes, specifically searching for genes annotated as aryl-sulfate sulfotransferases. Interestingly, we discovered that *Sutterella*, a genus within the human gut microbiome, contains multiple genes annotated for these sulfotransferases.

*Sutterella* species have been isolated from human fecal samples (35–41) and implicated in various human health conditions, including ulcerative colitis(42), autism spectrum disorders (ASD)(43, 44), bacteremia(45), multiple sclerosis(46), and autoimmune based thyroid disease(47). Additionally, *Sutterella* is known to degrade immunoglobulin A (IgA), suggesting a pro-inflammatory role (48). Interestingly, in the context of ulcerative colitis, a decline in *Sutterella wadsworthensis* has been associated with drug-free remission (42). Given its widespread presence and links to several human health concerns, we decided to investigate and understand the presence of annotated *asst* genes and their products (ASST proteins) in the genome of *S. wadsworthensis*. Our bioinformatics study has revealed a predominant presence of aryl-sulfate sulfotransferase (ASST) enzymes within the genus *Sutterella*. These enzymes exhibit considerable sequence and structural homogeneity, albeit with enough variability that can have functional consequences. Additionally, while sharing similarities with prokaryotic ASSTs such as those in *E. coli*, *Sutterella*’s sulfotransferases appear to have diverged evolutionarily from the PAPS-dependent class of sulfotransferases.

## Results and Discussion

### Distribution of aryl-sulfate sulfotransferase (*asst*) genes in human gut microbiota

Aryl-sulfate sulfotransferases (ASSTs) are enzymes found extensively across various genera of gut microbiota. Fig 2 illustrates the prevalence of annotated *asst* genes within diverse gut microbial genera. The data depicts the presence of these genes across different species and strains within each genus. The analysis omits numerous *E. coli* strains due to their high abundance and to avoid skewing the data representation.

**Fig 2.**
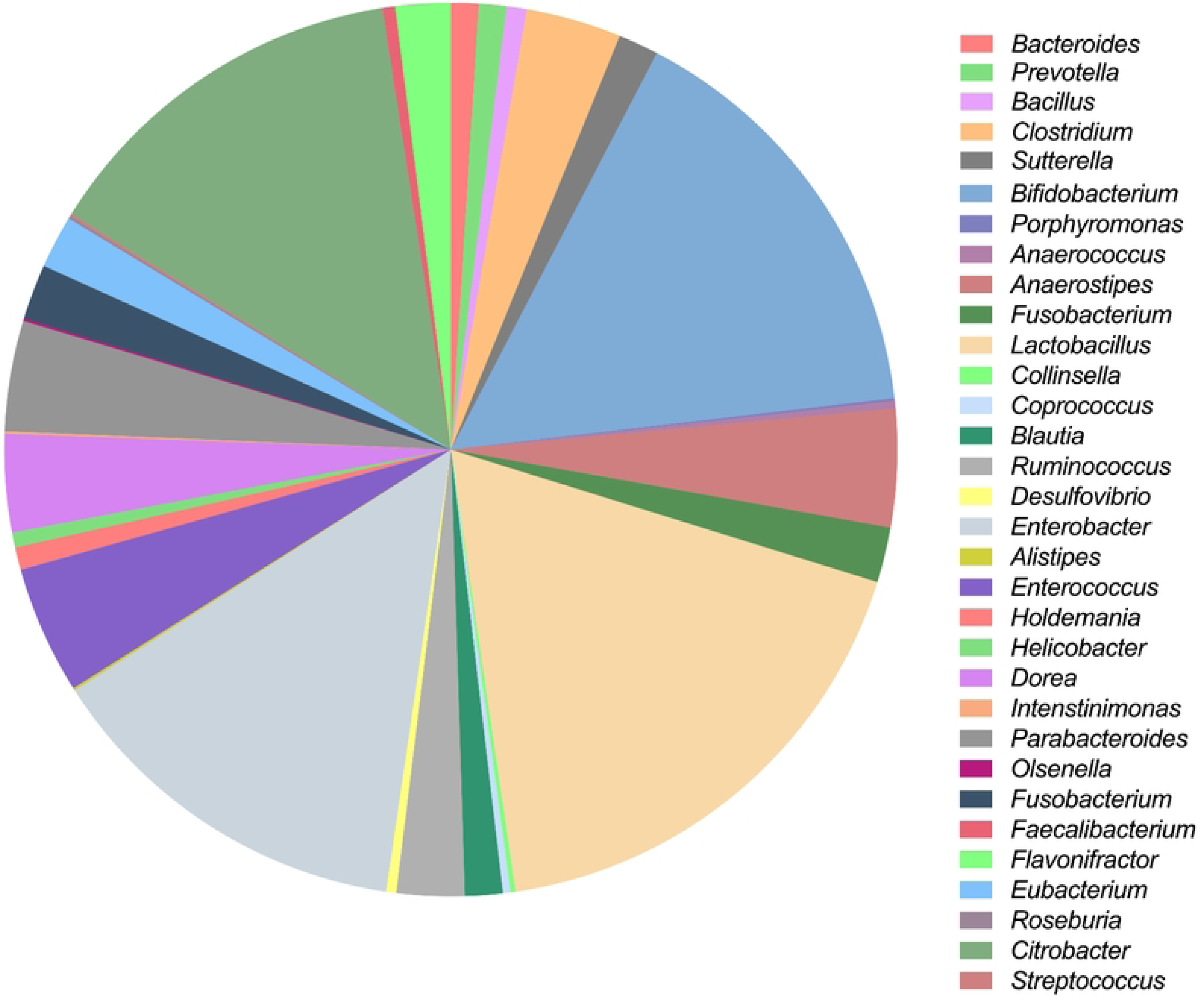
Distribution of annotated *asst* genes in the human gut microbes. The pie chart represents the distribution of annotated aryl-sulfate sulfotransferase (*asst*) genes across various known gut microbial genera. Each segment’s proportion reflects the relative count of *asst* genes within that particular genus.

Our research indicates a broad distribution of *asst* genes among gut microbes. Interestingly, a majority of these microbes possess only one or two annotated *asst* genes per organism. Genera such as *Lactobacillus*, *Bifidobacterium*, *Enterobacter*, and *Citrobacter* are predominant, collectively harboring 60% of all *asst* gene annotations identified in gut microbes. Despite the apparent ubiquity of these genes, the underlying reasons for their prevalence in human gut microbiota remain elusive.

### Variability of annotated aryl-sulfate sulfotransferase (*asst*) genes in the genus *Sutterella*

Our search through IMG genome database has uncovered multiple annotated aryl-sulfate sulfotransferases within the genus *Sutterella.* Bar graph in Fig 3 and S2 Table details the distribution of these genes among different species and strains of *Sutterella*. Notably, *Sutterella wadsworthensis* exhibits the largest number of annotated *asst* genes, with individual strains showing variability ranging from 8 to 17 genes.

**Fig 3.**
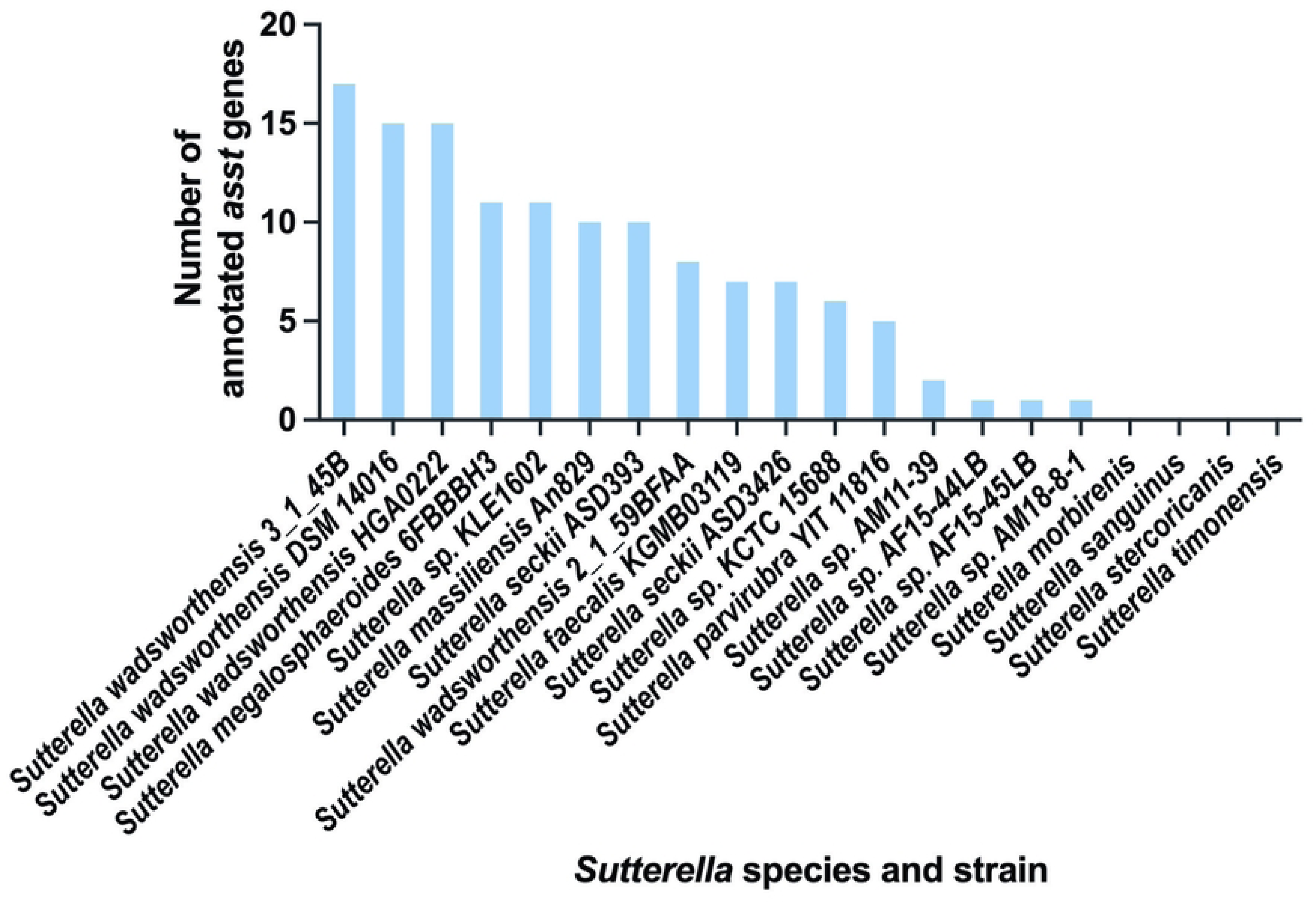
Annotated *asst* genes within the genus *Sutterella.* The bar graph represents the quantity of annotated *asst* genes across various species and strains of *Sutterella*, highlighting the substantial variability within the genus.

Furthermore, *Sutterella megalosphaeroides* and *Sutterella sp. KLE1602* each harbor a total of 11 annotated *asst* genes. Close behind, *Sutterella massiliensis* has 10 such genes. Distinct strains of *Sutterella seckii*, ASD393 and ASD3426, comprise 10 and 7 annotated *asst* genes, respectively. Other species including *Sutterella faecalis*, *Sutterella sp. KCTC 15688*, and *Sutterella parvirubra* display between 5 to 7 genes. Contrastingly, several strains such as AM11-39, AF15-44LB, AF15-45LB, and AM18-8-1 of *Sutterella sp.* contain only 1 to 2 annotated *asst* genes. However, species like *Sutterella morbirenis*, *Sutterella sanguinus*, *Sutterella stercoricanis*, and *Sutterella timonensis* have no detectable annotated *asst* genes.

This variation within the same genus raises intriguing questions: Why do certain *Sutterella* species have a multitude of *asst* genes while others have none? Addressing these questions will be possible only after the development of advanced genetic manipulation tools specifically tailored for the *Sutterella* genus.

### Bioinformatic analysis of annotated *asst* genes in *S. wadsworthensis 3_1_45B*

Given that *S. wadsworthensis* has the highest count of annotated genes encoding aryl-sulfate sulfotransferases (*assts*), we focused on a detailed characterization of these genes and their corresponding enzyme products from the strain *S. wadsworthensis 3_1_45B*, which contains 17 identified *asst* genes. The comprehensive list and characteristics of these 17 genes and enzymes can be found in S3 Table. For *asst* genes, we examined the percent GC content and the gene neighborhoods.

#### GC content variation in *asst* genes

We first analyzed the percent GC content of the *asst* genes from *S. wadsworthensis 3_1_45B*, with data sourced from the IMG genome database. By plotting the GC content, we aimed to determine any variations or similarities that might exist within *asst* genes. The GC content of various *S. wadsworthensis* strains is generally around 55%(45). The mean GC content of the *asst* genes was found to be 55.3%, with a median of 55%, aligning closely with the previously reported values for a GC content of *S. wadsworthensis*(45). However, the GC content of the 17 annotated *asst* genes did display some diversity, ranging from a minimum of 49% to a maximum of 61% (Fig 4).

**Fig 4.**
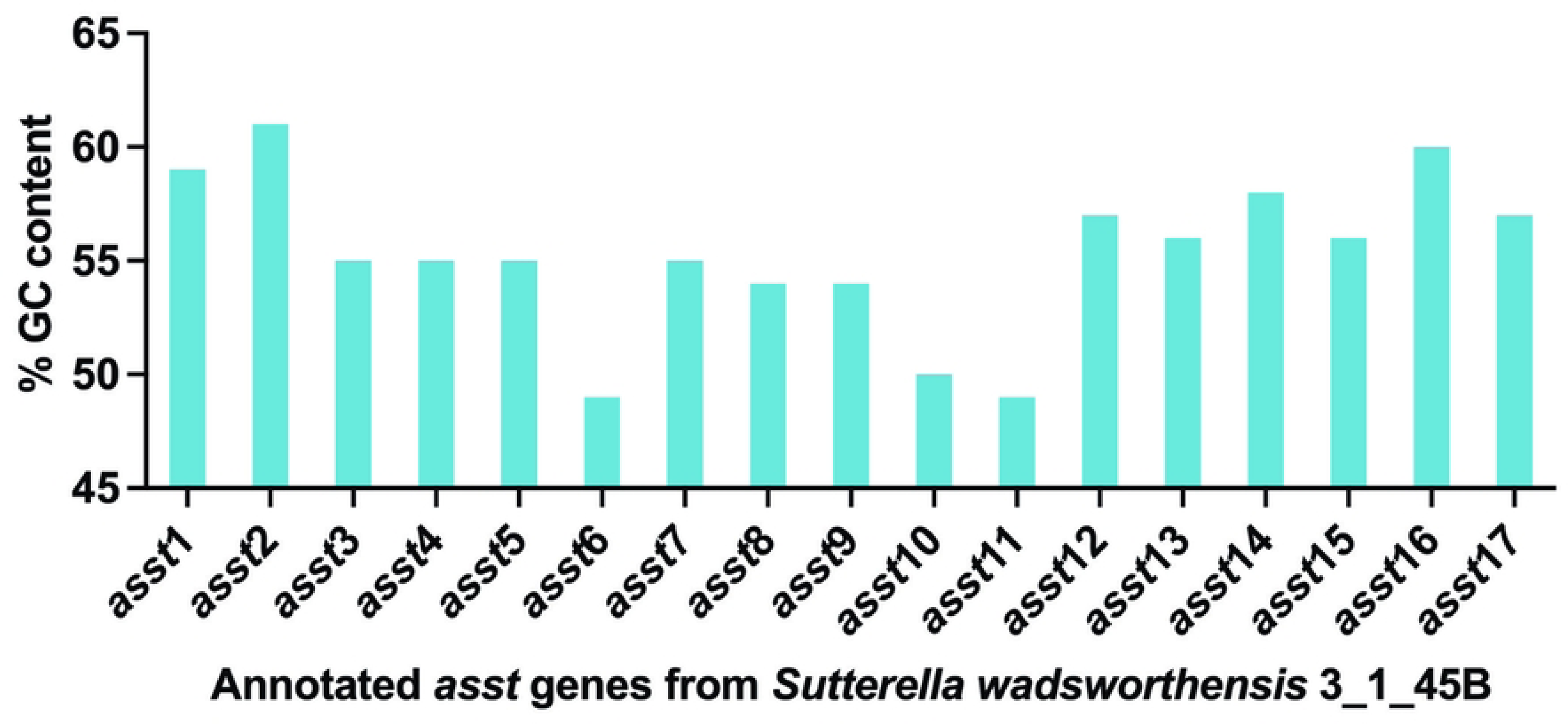
Variation in the %GC content of *asst* genes. This figure shows a bar graph representation of %GC content for all annotated *asst* genes from *S. wadsworthensis 3_1_45B*.

#### Analysis of gene neighborhoods for *asst* genes in *S. wadsworthensis 3_1_45B*

Investigating the gene neighborhood is critical for understanding the physiological roles of specific genes and identifying potential operon structures which may elucidate their functions. Interestingly, in *E. coli*, the *asst* gene is part of an operon associated with the formation of disulfide bonds (29). Accordingly, such analyses are important in shedding light on the functions of the annotated sulfotransferases. Fig 5 illustrates the comprehensive analysis of gene neighborhoods of *asst* genes in *S. wadsworthensis 3_1_45B*.

**Fig 5.**
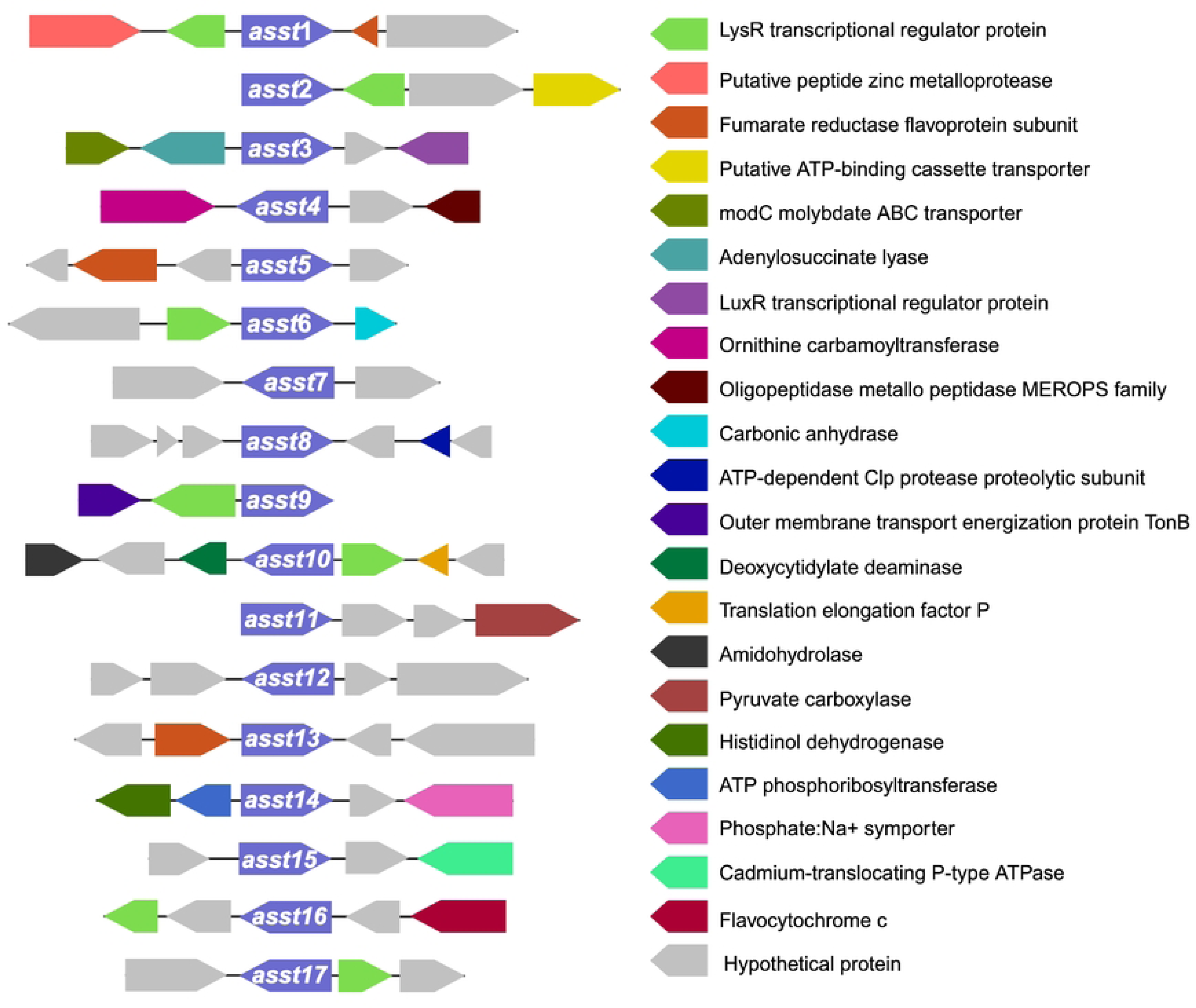
Gene neighborhood maps of annotated *asst* genes. Each gene is depicted as an arrow and is represented by a unique color. All *asst* genes (from 1-17) are represented in purple color. Size of each arrow is approximately proportional to the gene size.

An analysis of the seventeen annotated *asst* genes has revealed that seven of these have *lysR* gene in close vicinity (Fig 5 and Table 1). The *lysR* gene encodes LysR family transcription regulator. LysR-type transcriptional regulators (LTTRs) play dual roles as activators or repressors of gene expression(49). In *E. coli*, for instance, the LysR-type regulator CysB governs the expression of genes crucial for sulfate assimilation and organic sulfur utilization(50). This regulatory control extends to sulfate starvation responses in *P. putida* as well(51). Furthermore, recent discoveries have identified a novel LysR type transcriptional regulator (LTTR) from *A. baumannii* which is involved in controlling the expression of genes involved in the uptake and reduction of various sulfur compounds(52). LTTRs are prolific within prokaryotes and are known to regulate multitude of bacterial functions, such as stress response, antibiotic resistance, motility, quorum sensing, degradation of aromatic compounds, and biosynthesis of amino acids(53).

**Table 1.** List of genes found in the neighborhood of annotated *asst* genes that encode for *Sw*ASST proteins.

| Annotated Genes in the neighborhood of *asst* genes | Number of occurrences around *asst* genes | Found in the neighborhood of this *asst* gene |
| --- | --- | --- |
| LysR transcriptional regulator protein | 7 | 1, 2, 6, 9, 10, 16, 17 |
| Fumarate reductase flavoprotein subunit | 3 | 1, 5, 13 |
| Putative peptide zinc metalloprotease | 1 | 1 |
| Putative ATP-binding cassette transporter | 1 | 2 |
| LuxR transcriptional regulator protein | 1 | 3 |
| modC molybdate ABC transporter | 1 | 3 |
| Adenylosuccinate lyase | 1 | 3 |
| Ornithine carbamoyltransferase | 1 | 4 |
| Oligopeptidase metallo peptidase MEROPS family | 1 | 4 |
| Carbonic anhydrase | 1 | 6 |
| ATP-dependent Clp protease proteolytic subunit | 1 | 8 |
| Outer membrane transport energization protein TonB | 1 | 9 |
| Translation elongation factor P | 1 | 10 |
| Deoxycytidylate deaminase | 1 | 10 |
| Amidohydrolase | 1 | 10 |
| Pyruvate carboxylase | 1 | 11 |
| ATP phosphoribosyltransferase | 1 | 14 |

Information from Fig 5 shown as a list.

Given that the ASST proteins are predicated to catalyze sulfate transfer reactions, it is plausible that LysR may influence the expression of ASSTs. Our findings indicate that *lysR* is a predominant gene in the vicinity of genes encoding *Sw*ASST proteins. Other genes located near *assts* are listed in Table 1, providing a comprehensive view of the gene neighborhood.

In the vicinity of *asst* genes, particularly those encoding *Sw*ASSTs 1, 5, and 13, we frequently identified the gene for the fumarate reductase flavoprotein subunit. This protein is integral to the process of anaerobic respiration(54), where sulfate can serve as one of the terminal electron acceptors. In addition, certain gut microbes are known to use naturally occurring organosulfonates, such as taurine, as terminal electron acceptors(55). Given that aryl-sulfate sulfotransferases (ASSTs) are involved in the production of organosulfonates through the transfer of sulfates between organic molecules(23, 24, 26), it raises the question of whether fumarate reductase could facilitate the transfer of electrons to these organosulfonates. Could there be a coordinated expression between ASSTs and fumarate reductase, suggesting a metabolic linkage? This presents an interesting hypothesis which requires experimental validation. Beyond the two genes commonly found near various *asst* genes, our analysis has identified additional genes situated in close proximity to single and specific *asst* genes encoding *Sw*ASSTs.

For instance, in the gene neighborhood of *asst1*, there is a gene that encodes a putative zinc metalloprotease. These enzymes are ubiquitous and versatile, often associated with pathogenicity and virulence in pathogenic microbes, though they are also present in non-pathogenic species. The presence of zinc metalloprotease is speculated to confer an evolutionary advantage to the microorganisms that retains it(56). However, the connection between an ASST protein and zinc metalloprotease is unclear.

In the vicinity of the *asst2* gene, there is a gene that encodes a putative ATP-binding cassette transporter (ABC transporter). ABC transporters, present in both eukaryotes and prokaryotes, perform a wide range of functions. Among their roles in microbial organisms is the transportation of sulfate, sulfonates, and sulfate esters(57, 58). Further research may reveal whether this ABC transporter is responsible for the translocation of the products generated by the activity of *Sw*ASST2.

Similar to the LysR family, the LuxR family transcriptional regulators can also function as either activators or repressors of gene expression, predominantly managing genes associated with quorum sensing. However, their regulatory influence extends beyond quorum sensing to other critical microbial biological functions(59). One such case is a novel LuxR-type regulator involved in catechol metabolism within the prominent gut microbe, *E. lenta*(60). A gene for LuxR family transcriptional regulator is also found adjacent to the *asst3* gene which encodes *Sw*ASST3. Whether it serves as a regulator for the expression of the *asst* gene, potentially influencing the metabolism of sulfated molecules in *Sutterella* spp., is a question that remains open for investigation.

Adjacent to the *asst3* encoding *Sw*ASST3 are two other genes in addition to the gene that produces LuxR regulator (Table 1). The first is an annotated gene which encodes molybdate ABC transporter. Previous studies have demonstrated the competitive interaction between sulfate ions and molybdate for uptake via similar transport systems(61, 62), an observation that has been made in the small intestine of sheep(63). This raises the question: might there be a similar competitive mechanism affecting the transport of the sulfated products created by *Sw*ASST3 and molybdate?

The other gene in proximity to *asst3* encodes adenylosuccinate lyase, an enzyme implicated in purine biosynthesis via the de novo pathway. Its enzymatic activity is essential for the synthesis of AMP(64). Any possible connection between an ASST protein and an adenylosuccinate lyase has not been explored before.

Surrounding the gene encoding *Sw*ASST4, two notable genes are present (Table 1). The gene for ornithine carbamoyltransferase, which modulates ornithine levels, is one such gene. Microbes capable of breaking down ornithine demonstrate a competitive advantage(65). This raises the question: could this enzyme also confer a survival benefit to *Sutterella wadsworthensis* in a healthy gut, and might *Sw*ASST4 play a role in this dynamic? Further investigation is needed to explore these possibilities. Additional gene adjacent to *asst4* is a gene for oligopeptidase A, part of the M03A family of metallopeptidases, known for their ability to cleave small peptides starting with alanine or glycine(66). There are no reports on any interaction between an oligopeptidase A and an ASST.

In the vicinity of the gene (*asst6*) encoding *Sw*ASST6, lies a gene for a LysR family transcription regulator, previously discussed (Table 1). Another neighboring gene encodes carbonic anhydrase, pivotal for maintaining CO_2_, HCO ^-^, and H^+^ balance within microbes and facilitating crucial exchange of these molecules with various metabolic pathways. Gut microbial carbonic anhydrases significantly differ from those in the host(67), suggesting that their activity levels can impact the survival of gut microbiota(68). The gene neighborhood of *asst8* includes an ATP-dependent Clp protease gene (Table 1). Proteases like these are essential for the removal of damaged or misfolded proteins. Because of their critical role in protein degradation pathways, these proteins are imperative for microbial physiology(69).

The gene encoding *Sw*ASST9 is associated with two distinct genes within its local genomic landscape (Table 1). Besides the previously noted LysR family transcription regulator, this neighborhood also includes a gene for the outer membrane transport protein, TonB. TonB-dependent transporters are known to leverage the proton motive force to enable the translocation of substances across the outer membrane. Specifically, TonB plays a critical role in importing nutrients, particularly large polysaccharides derived from the diet, into gut microbes. The considerable size of these molecules necessitates an energy-dependent mechanism for membrane passage, a process facilitated by TonB-dependent transporters(70, 71).

In addition to an above mentioned *lysR* family transcription regulator gene, three other annotated genes surrounding *asst10* gene that encodes *Sw*ASST10 (Table 1). These genes are for translation elongation factor P, deoxycytidylate deaminase, and an amidohydrolase. The translation elongation factor P is instrumental in microbial protein synthesis, particularly for polypeptides comprising polyproline sequences(72). While the deoxycytidylate deaminase plays a crucial role in the synthesis of the thymidine nucleotide, a building block of DNA (73). The amidohydrolase, a versatile enzyme, is capable of cleaving a variety of bonds such as carbon-oxygen, phosphorous-oxygen, phosphorous-sulfur, carbon-nitrogen, carbon-sulfur, and carbon-chlorine, with known involvement in the metabolism of xenobiotics(74).

Adjacent to the gene encoding *Sw*ASST11, pyruvate carboxylase has been identified, as listed in Table 1. This enzyme plays a pivotal role in anaplerosis, the process of replenishing intermediates of the tricarboxylic acid (TCA) cycle that are consumed by various biosynthetic pathways. Pyruvate carboxylase specifically catalyzes the formation of oxaloacetate, a key intermediate in the TCA cycle. Moreover, carboxylases are integral to carbon fixation reactions, facilitating the incorporation of inorganic carbon into organic molecules (75).

Surrounding the gene encoding *Sw*ASST14, three significant genes have been annotated: histidinol dehydrogenase, ATP phosphoribosyltransferase, and a phosphate:Na+ symporter, as noted in Table 1 and Fig 5. The biosynthesis of histidine is a well-conserved pathway found across various life forms, with the exception of mammals(76). ATP phosphoribosyltransferase and histidinol dehydrogenase are key enzymes in this pathway, representing the first and the final steps, respectively(77). ATP phosphoribosyltransferase initiates the pathway by condensing 1-(5-phospho-D-ribosyl)-ATP with a diphosphate molecule. The phosphate:Na+ symporter may facilitate the uptake of phosphate, the substrate required for this first reaction in histidine biosynthesis. The potential involvement of *Sw*ASST14 in histidine biosynthesis or its regulation poses an intriguing question for future research endeavors.

Located in the genomic vicinity of the gene encoding *Sw*ASST15 is a cadmium-translocating P-type ATPase (Table 1). P-type ATPases play a critical role in cellular homeostasis by regulating the intracellular concentrations of both essential and toxic transition metals. They achieve this by actively expelling toxic metals from the cells, thereby mitigating metal toxicity(78).

Lastly, there are two annotated genes adjacent to the gene encoding *Sw*ASST16 (Table 1). One is the previously mentioned LysR family transcription regulator, and the other encodes flavocytochrome c. Flavocytochromes c are typically produced in large quantities under anaerobic conditions(79). They are prominent in sulfate-reducing microbes, where soluble flavocytochrome c proteins transfer electrons to various acceptors, including sulfate, thiosulfate, sulfur, nitrate, and fumarate(80). The involvement of flavocytochrome c proteins in the reduction of organosulfates has not been established in the literature. For this reason, the likelihood of *Sw*ASST16 reaction products which are organosulfates, acting as direct terminal electron acceptors for flavocytochrome c seems minimal. Nonetheless, it would be intriguing to explore whether these organosulfates participate indirectly in a broader metabolic network that includes both ASST and flavocytochrome c proteins, thereby contributing to the electron transfer to organosulfates through more complex pathways.

Our gene neighborhood analysis has indicated that asst genes from *S. wadsworthensis 3_1_45B* typically do not integrate into recognizable operons. Instead, they appear to exist independently within the genome, as depicted in Fig 5. This finding leads us to hypothesize that the *asst* genes generally function as isolated units rather than as part of operonic clusters. Nevertheless, to fully understand the biological relevance of this arrangement, experimental investigations into the interactions between *Sw*ASSTs and the products of the neighboring genes are warranted.

### Bioinformatic analysis of predicted ASST proteins from *S. wadsworthensis 3_1_45B*

For the predicted *Sw*ASST proteins, we assessed protein length, presence of signal peptides, probable cellular localization, presence of transmembrane regions, amino acid sequence conservation, and structural variability.

#### Protein length variation among ASST enzymes of *S. wadsworthensis 3_1_45B*

The data presented in Fig 6 and S3 Table, details the amino acid lengths of the 17 identified ASST enzymes from *S. wadsworthensis 3_1_45B*. The majority of these enzymes consist of approximately 600 amino acids. Specifically, the longest ASST comprises 615 amino acids, while the shortest contains 449 amino acids. 16 of the 17 ASST proteins have lengths within a narrow range of 605 to 615 amino acids, exhibiting consistency in size among these enzymes.

**Fig 6.**
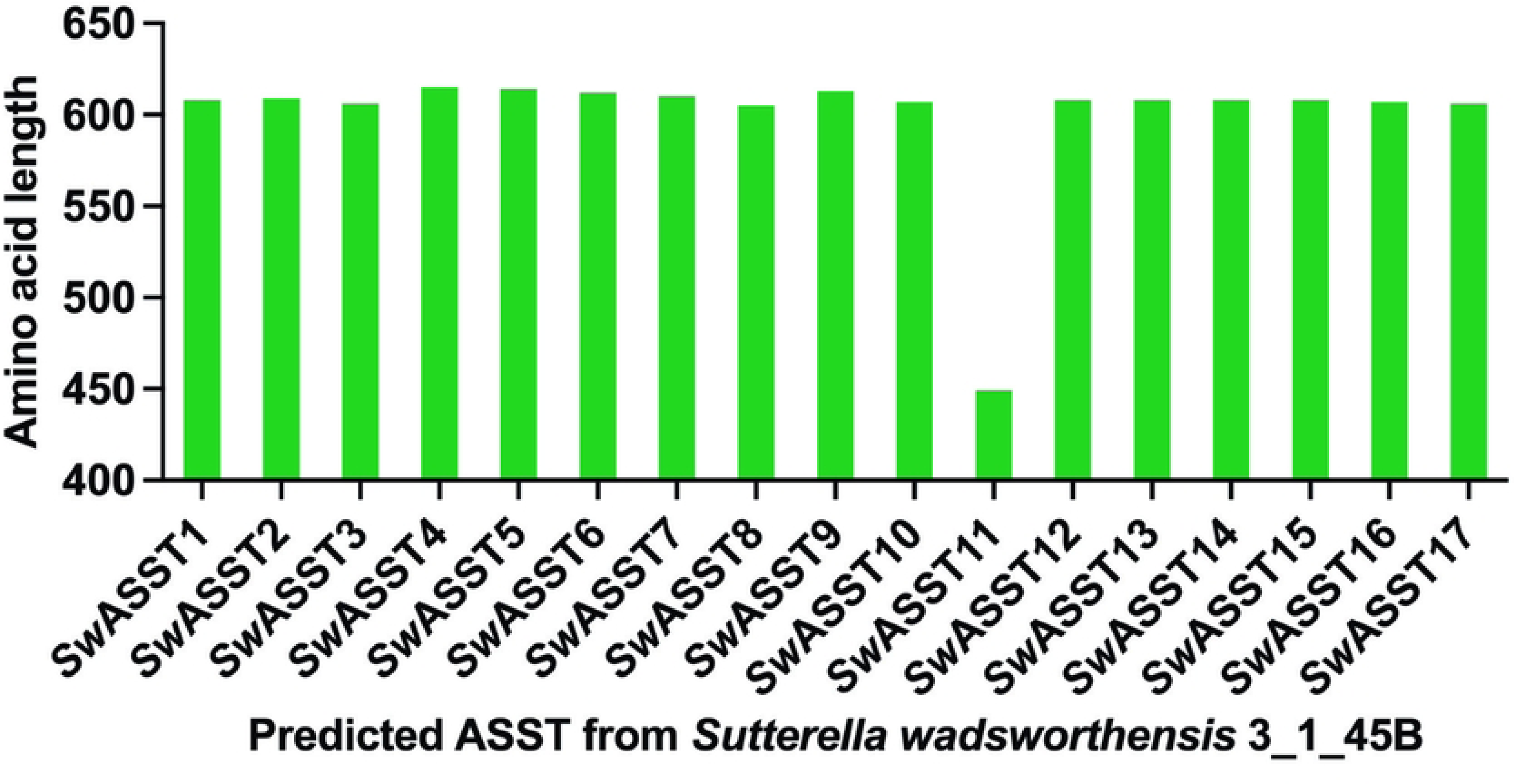
Variation in the length of ASST enzymes of *S. wadsworthensis 3_1_45B*. The plot illustrates the range of amino acid counts as comparison of protein lengths across the 17 predicted ASST enzymes.

#### Analysis of signal peptides in *Sw*ASST enzymes

Given that a well-characterized ASST from *E. coli* has a signal peptide(29), we conducted a sequence analysis to determine the prevalence and types of signal peptides in *Sw*ASSTs from *S. wadsworthensis 3_1_45B*. Our findings reveal that 16 of the 17 annotated *Sw*ASSTs are predicted to have signal peptides, suggesting their translocation across the cytoplasmic membrane (Fig 7A and S3 Table). The analysis of these signal peptides was carried out using SignalP 6.0, confirming the annotations provided by the IMG genome database. Upon examining the lengths of these signal peptides, the average length was found to be 26 amino acids, with a median of 25.5 amino acids. Among the 16 *Sw*ASSTs with signal peptides, 10 (*Sw*ASSTs 2, 4, 7, 8, 9, 10, 12, 13, 15, 17) harbor the Sec-type signal peptides associated with the general secretion pathway, while 6 (*Sw*ASSTs 1, 3, 5, 6, 14, 16) contain the Tat-type signal peptides linked to the twin-arginine translocation pathway (Fig 7B and Table 2). In addition, there is significant variability in the amino acid sequences of both Sec and Tat type signal peptides across the different annotated *Sw*ASSTs (Fig 8A and 8B).

**Fig 7.**
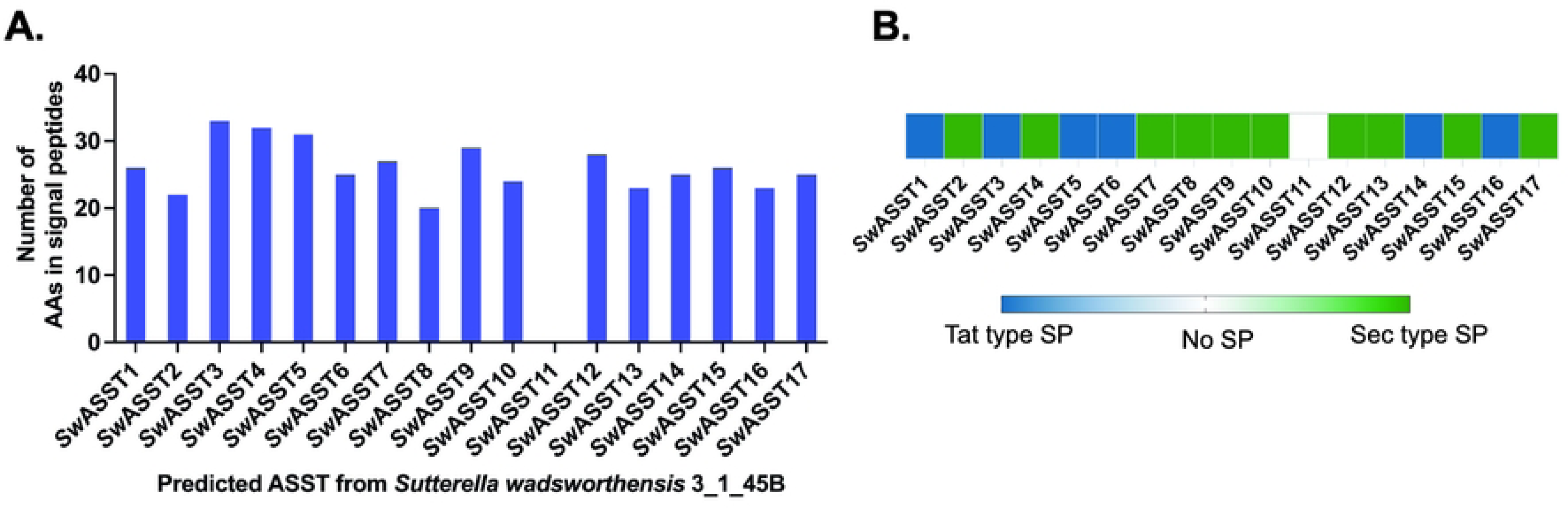
Signal peptide characteristics in predicted *Sw*ASSTs. A) Depicts the length of signal peptides in amino acid (AA) residues. B) Categorizes the signal peptides by type (Sec or Tat) within the *Sw*ASSTs, where Sec type is represented as green and Tat type as blue.

**Fig 8.**
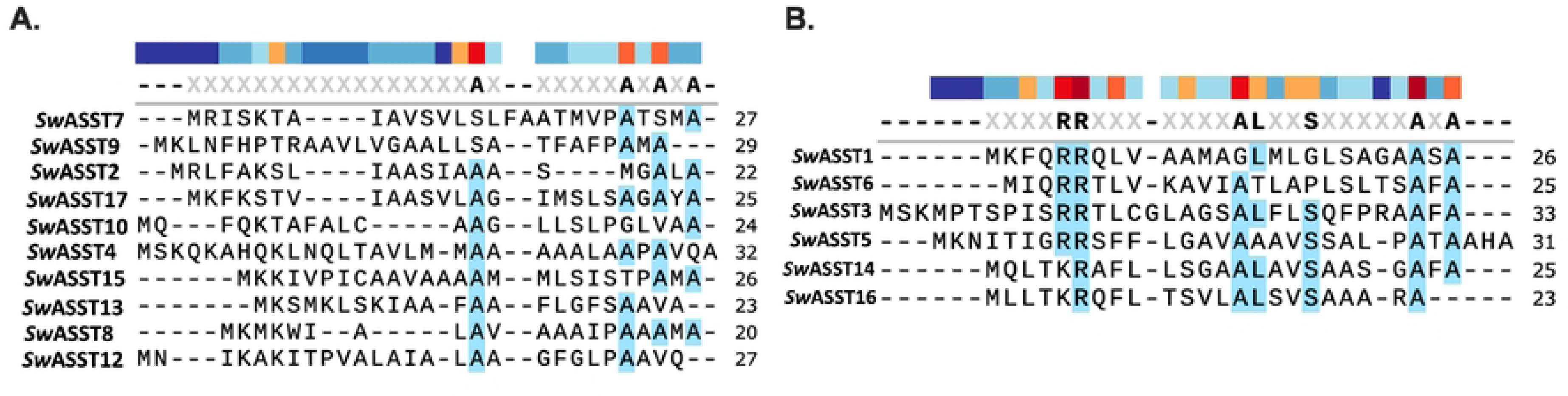
Sequence alignments for signal peptides of *Sw*ASSTs. A) Sequence alignment for Sec type signal peptides of *Sw*ASSTs. B) Sequence alignment for Tat type signal peptides of *Sw*ASSTs. Color bar above the sequences shows the amino acid conservation at that position, where dark red is highly conserved and dark blue indicates the least conserved position.

**Table 2.**
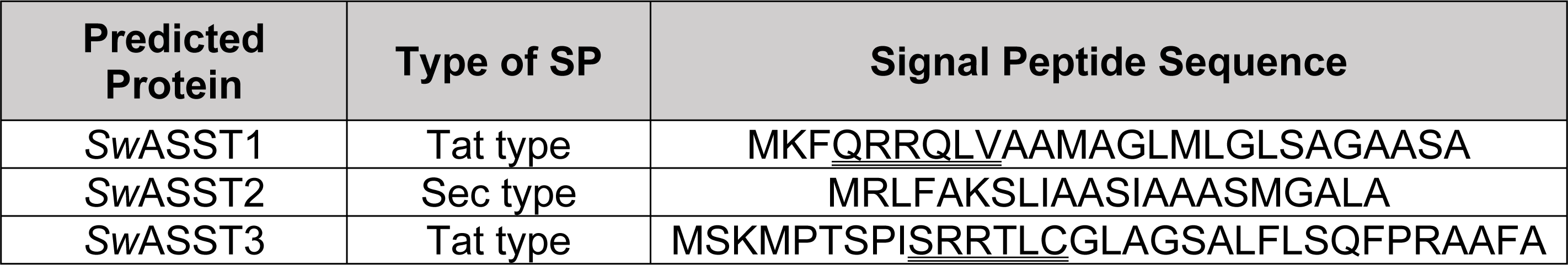
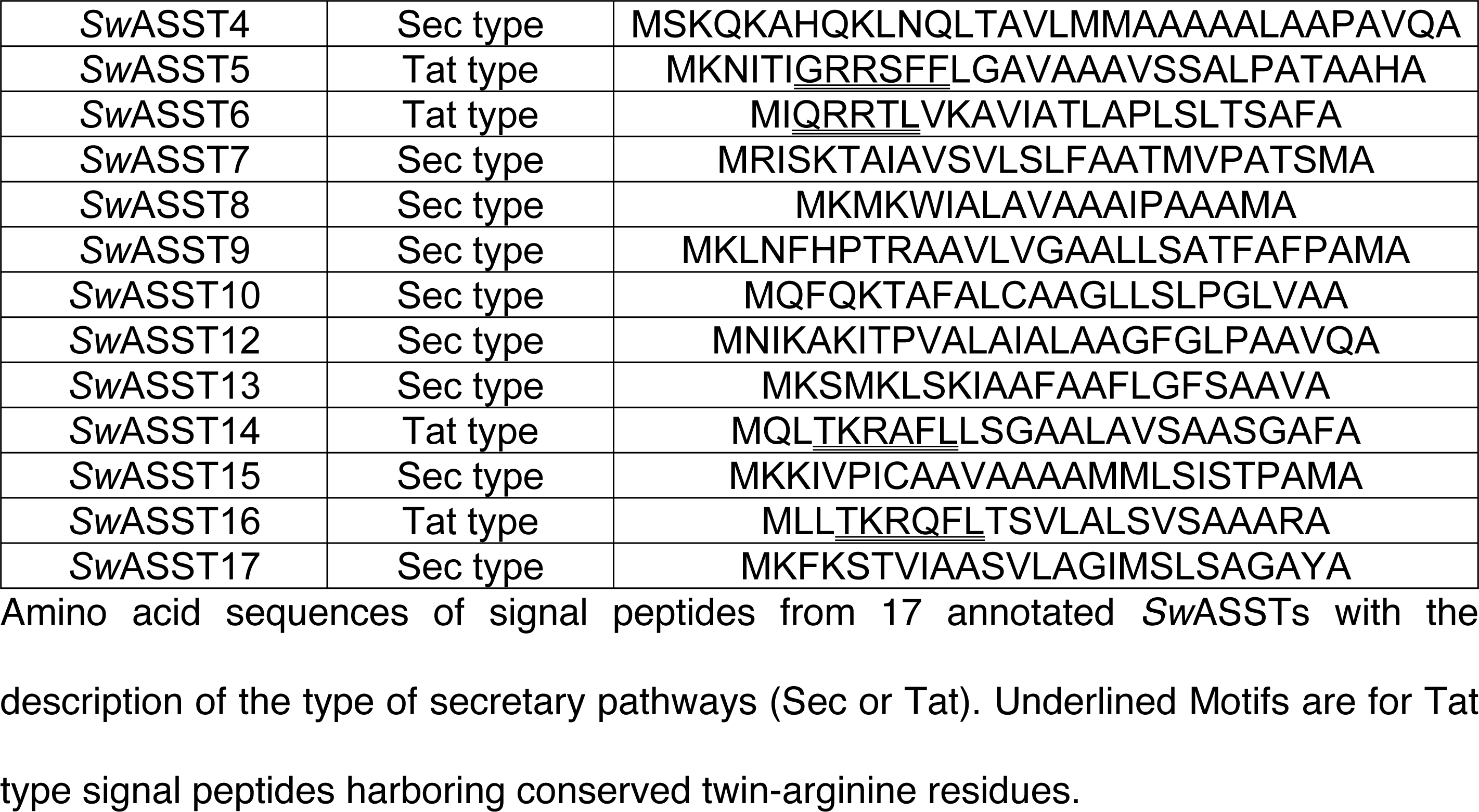
Signal peptide sequences and types of *Sw*ASSTs.

Sec and Tat translocases are common mechanisms for the transfer of proteins across the cytoplasmic membrane. The Sec system mediates protein transport across the membrane in an unfolded state, whereas the Tat system transports proteins in a folded state(81). These proteins have distinct N-terminal signal peptides, with the Sec pathway typically featuring a positively charged N-terminus, a hydrophobic core, and polar C-terminal residues (Fig 8A and Table 2). The Tat pathway signal peptides are more conserved, particularly the twin arginine residues (Fig 8B). However, *Sw*ASST14 and *Sw*ASST16 diverge from this pattern, presenting a lysine (K) in place of the first arginine (R) in the conserved region—a rare variant with potential implications for translocation efficiency, as reported in literature(82, 83). This observation suggests that *Sw*ASST14 and 16 may have a lower translocation efficiency than other *Sw*ASSTs with canonical twin-arginine motifs. Additionally, both Sec and Tat signal peptides frequently contain the conserved A-X-A motif at the C-terminus, a feature commonly found in N-terminal signal peptides(84).

#### Analysis of Transmembrane Domains in *Sw*ASST Proteins from *S. wadsworthensis 3_1_45B*

Preliminary investigations using the IMG genome database indicated that transmembrane (TM) helices might exist in certain annotated *Sw*ASST proteins. To verify and further understand the structural architecture of these proteins, we employed the DeepTMHMM tool for a detailed analysis of their TM helices (85, 86). Our analysis confirmed that 9 of the 17 *Sw*ASSTs have predicted TM regions (Fig 9 and S4 Table).

**Fig 9.**
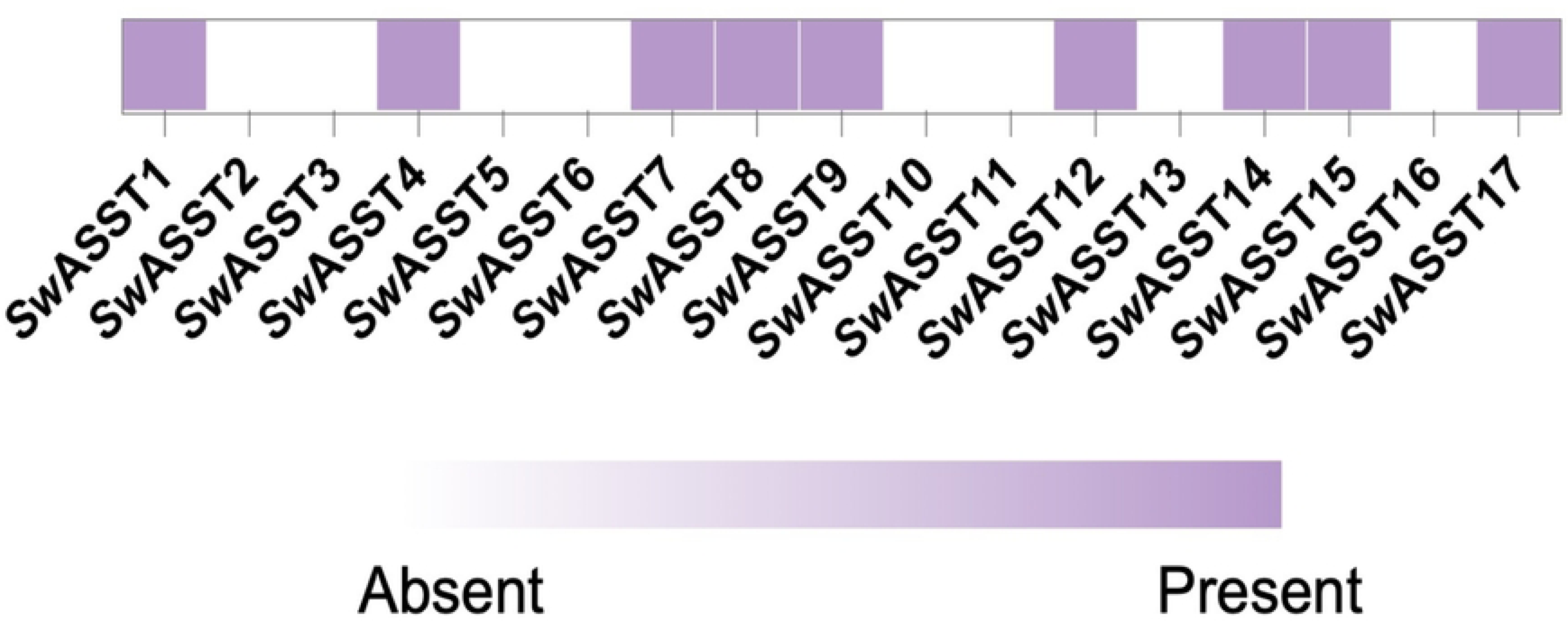
Transmembrane regions of *Sw*ASSTs. The heat map depicts the presence (purple) or absence (white) of predicted transmembrane helices in *Sw*ASSTs.

As previously mentioned, 16 of the 17 *Sw*ASSTs are characterized by Sec or Tat signal peptides at their N-terminus. Proteins secreted through the Sec or Tat pathways may localize to the periplasm, integrate into the cytoplasmic membrane, or be exported outside the cell via other secretion systems, especially in Gram-negative bacteria(87).

Periplasmic and extracellular proteins typically follow a SecB-mediated route, with extracellular proteins requiring an additional translocation step across the outer membrane via Type II or Type V secretion systems (T2SS or T5SS)(87). Genomic analysis of *S. wadsworthensis 3_1_45B* revealed the presence of T2SS components, suggesting that some *Sw*ASSTs might be T2SS substrates.

Additionally, proteins secreted by the Sec pathway that embed in the inner cytoplasmic membrane utilize an SRP-mediated pathway, which requires the presence of TM regions within the protein substrates. Table 3 combines our findings regarding *Sw*ASSTs harboring the Sec or Tat signal peptides, delineating those with and without TM domains, and proposes their potential cellular destinations.

**Table 3.**
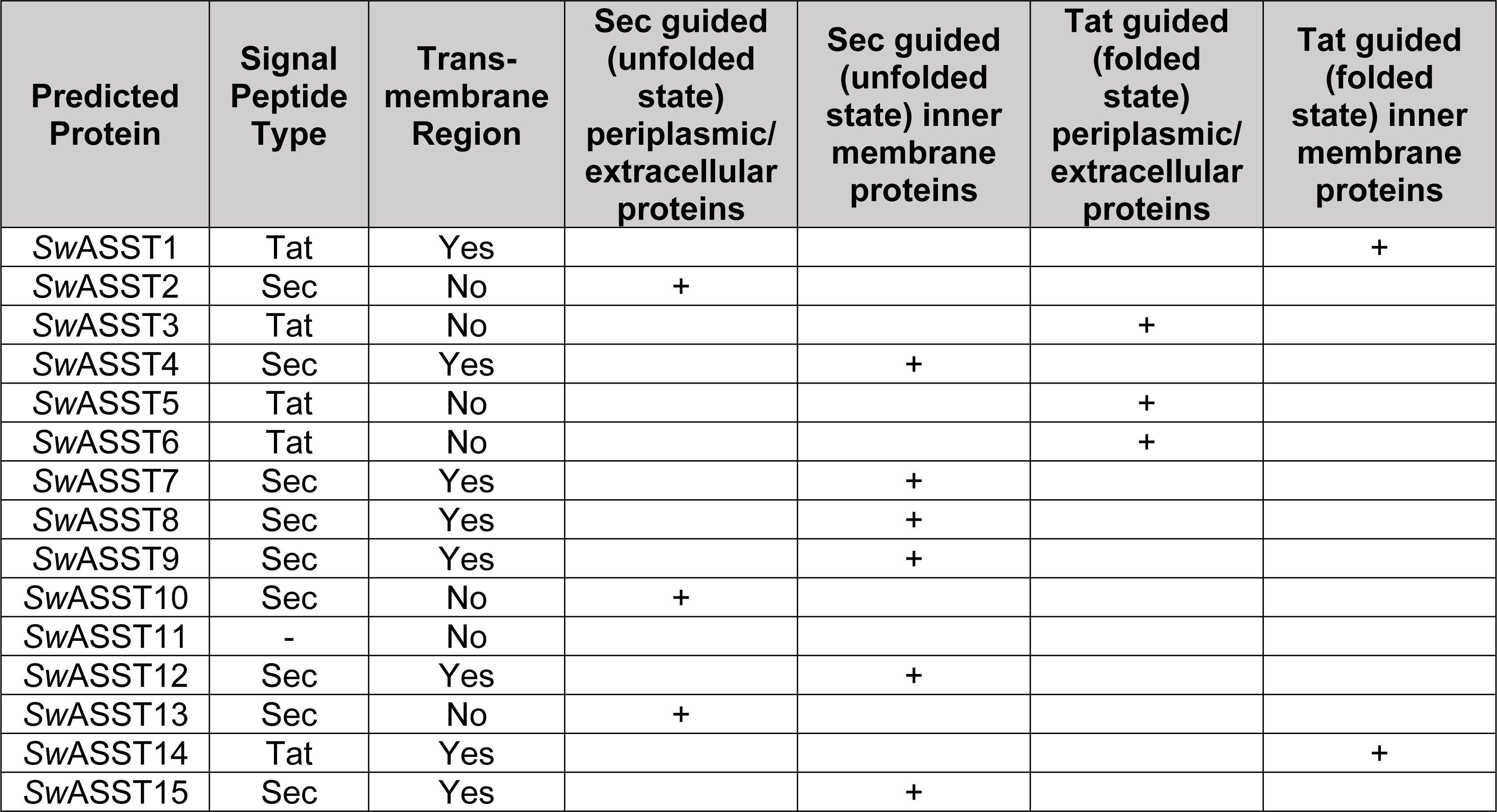
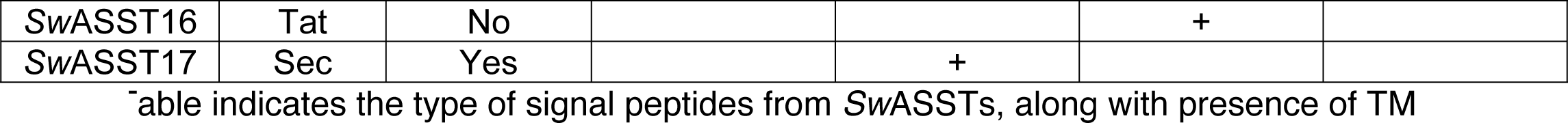
Potential cellular destinations of *Sw*ASSTs.

| Predicted Protein | Signal Peptide Type | Trans-membrane Region | Sec guided (unfolded state) periplasmic/extracellular proteins | Sec guided (unfolded state) inner membrane proteins | Tat guided (folded state) periplasmic/extracellular proteins | Tat guided (folded state) inner membrane proteins |
| --- | --- | --- | --- | --- | --- | --- |
| SwASST1 | Tat | Yes |  |  |  | + |
| SwASST2 | Sec | No | + |  |  |  |
| SwASST3 | Tat | No |  |  | + |  |
| SwASST4 | Sec | Yes |  | + |  |  |
| SwASST5 | Tat | No |  |  | + |  |
| SwASST6 | Tat | No |  |  | + |  |
| SwASST7 | Sec | Yes |  | + |  |  |
| SwASST8 | Sec | Yes |  | + |  |  |
| SwASST9 | Sec | Yes |  | + |  |  |
| SwASST10 | Sec | No | + |  |  |  |
| SwASST11 | - | No |  |  |  |  |
| SwASST12 | Sec | Yes |  | + |  |  |
| SwASST13 | Sec | No | + |  |  |  |
| SwASST14 | Tat | Yes |  |  |  | + |
| SwASST15 | Sec | Yes |  | + |  |  |
| SwASST16 | Tat | No |  |  | + |  |
| SwASST17 | Sec | Yes |  | + |  |  |

Table indicates the type of signal peptides from *SwASSTs*, along with presence of TM

#### Sequences similarity analysis of *Sw*ASSTs

For the multiple sequence alignments, *E. coli* ASST (*Ec*ASST) served as a reference sequence because it has been studied well in the context of structure-function relationship(29). Utilizing the Clustal Omega tool from UniProt for multiple sequence alignment, we observed that all the key active site residues of *Ec*ASST, crucial for catalysis are all well conserved across all 17 *Sw*ASSTs. Especially, residues corresponding to His-252 (H252), His-356 (H356), Asn-358 (N358), Arg-374 (R374), and His-436 (H436)(29) (Fig 10). His-436 is the most important catalytic residue that gets transiently sulfated during the catalysis by *E. coli* sulfotransferase. This happens due to two half reactions that occur during the ASST catalysis. In the first half reaction, a donor adds a sulfate group to the active site histidine residue. In the second half reaction, this sulfate group from a histidine residue is transferred to the acceptor molecule which completes the catalytic cycle. The multiple sequence alignment of all 17 *Sw*ASST enzymes confirms that this catalytic histidine residue is perfectly conserved, suggesting a similar catalytic mechanism may be at play within these enzymes (Fig 10).

**Fig 10.**
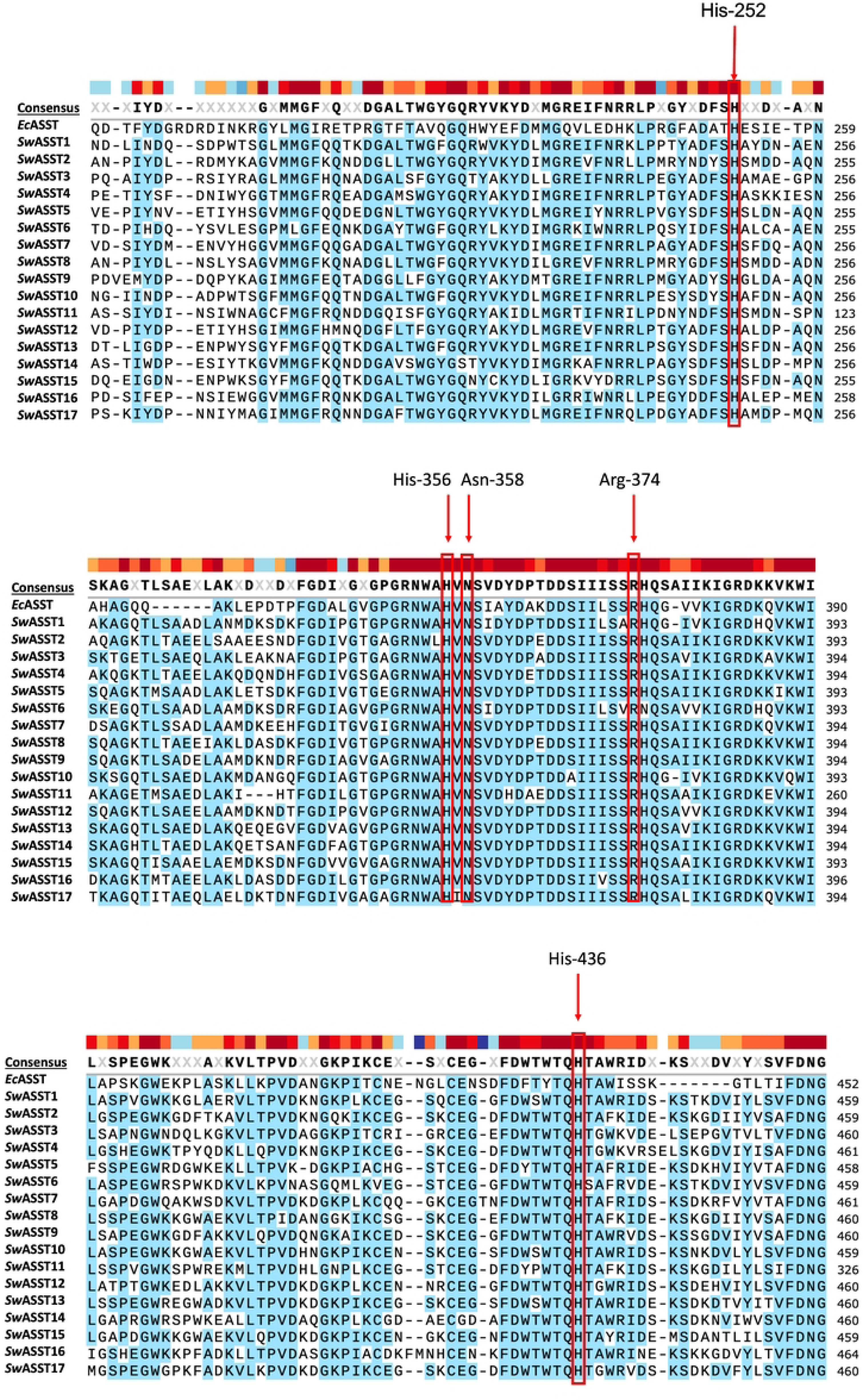
Multiple sequence alignment of *Sw*ASSTs. Alignment of *Sw*ASSTs with *Ec*ASST highlighting catalytically crucial residues in red. Color bar above the consensus line shows the amino acid conservation at that position. Dark red is highly conserved, while dark blue is little to no conservation.

This alignment revealed an average sequence similarity of 62% among all 17 *Sw*ASSTs. The range of similarity spanned from a minimum of 53% to a maximum of 75.5%, with the median and mode being 62% and 61.25%, respectively (Fig 11A). The degree of similarity indicates that while the *Sw*ASST enzymes share a common structural framework, there is sufficient variation to suggest they may have specialized functions. Interestingly, *Sw*ASST2 and *Sw*ASST8 exhibited the greatest sequence similarity at 75.5%, whereas *Sw*ASST1 and *Sw*ASST3 shared the least similarity at 53%.

**Fig 11.**
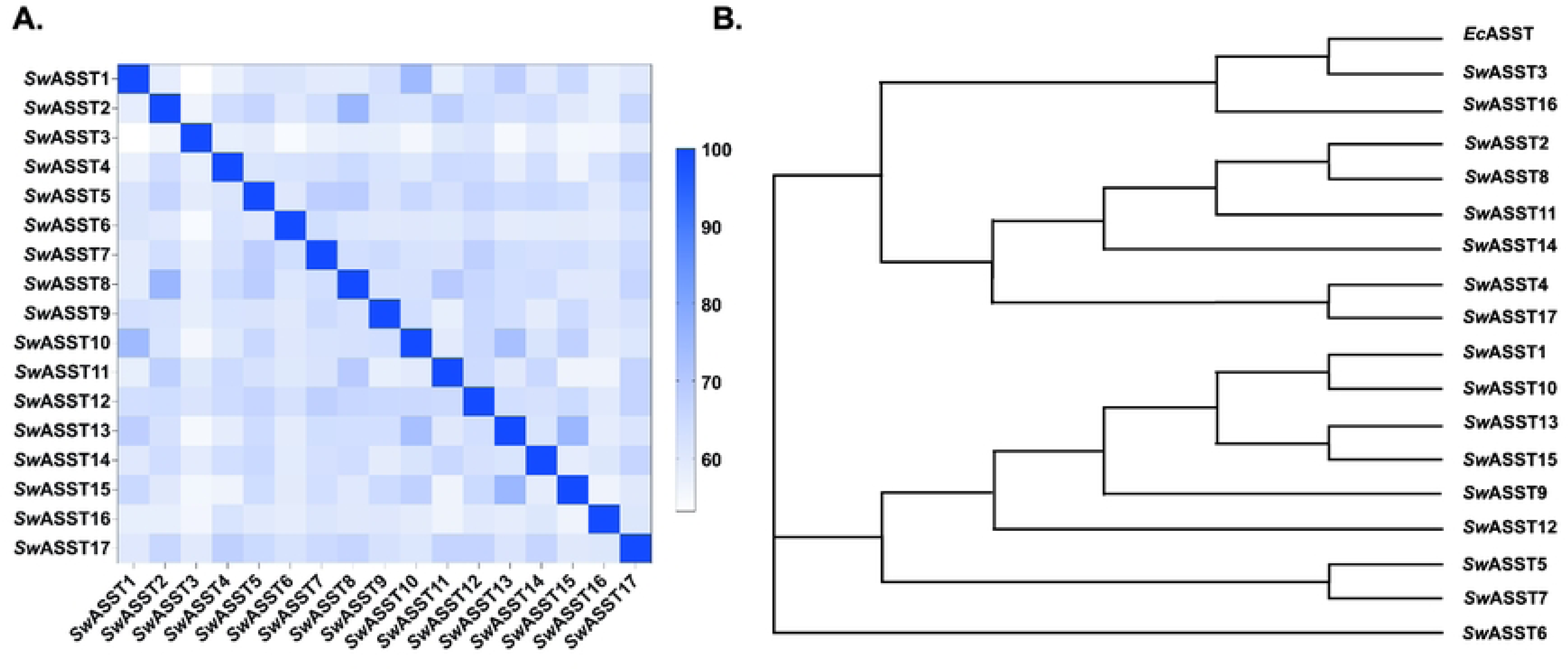
Sequence similarity analysis of *Sw*ASSTs from *S. wadsworthensis 3_1_45B.* A) Percent identity matrix B) Phylogenetic tree of the 17 predicted *Sw*ASSTs and *E. coli* ASST (*Ec*ASST) from *E. coli* strain CFT073.

Additionally, our sequence alignments show that *Sw*ASSTs harbor more sequence similarities with each other than with *Ec*ASST. There are residues that are absolutely conserved in *Sw*ASSTs but are completely different in *Ec*ASST (S1 Fig). For example, there are two conserved asparagine residues in all *Sw*ASSTs (1^st^ one around 160-163 in and 2^nd^ one around 161-164 in all *Sw*ASSTs except in the shorter *Sw*ASST11 where these are located at positions 28 and 29) which are not conserved in *Ec*ASST. *Ec*ASST has histidine and glycine residues in this place. Similarly, a threonine (T166) is replaced by an absolutely conserved glycine (164–167) residue in all *Sw*ASSTs. There are more such examples which can be seen in extended sequence alignment of these proteins. Based on sequence alignment, it is apparent that *Sw*ASST3 and 16 are more closely related to *Ec*ASST protein (Fig 11B, phylogenetic tree).

#### Structural variations in *Sw*ASST proteins

To understand differences at the structure level, structures of all 17 *Sw*ASSTs were generated using Alphafold2. The resulting confidence levels of each model, measured by the local distance difference test (lDDT) score, are plotted against their respective amino acid positions which were generated by Alphafold2 (S2 Fig)(88). Generally, the lDDT confidence plots show decreased prediction reliability around residue positions 200, 350, and 550 for most *Sw*ASSTs. However, *Sw*ASST3, *Sw*ASST12, and *Sw*ASST16 exhibit less pronounced dips in confidence around residue 550. *Sw*ASST11, being smaller, displays a distinct confidence profile with no notable decrease in prediction reliability around the residues corresponding to position 550 in the other *Sw*ASSTs, such as in *Sw*ASSTs 3, 12, and 16. These observations highlight subtle structural differences among the *Sw*ASST proteins, possibly reflecting variations in their functional attributes. Structural alignment of *Sw*ASSTs, conducted using PyMol, revealed a shared common fold among these enzymes (Fig 12). The ability to superimpose these structures, shown in Fig 13, further supports their structural convergence. Additionally, an average root-mean-square deviation (RMSD) of 0.52 Å across all 17 aligned structures suggests a high degree of similarity(89). While the overall secondary structure elements, such as alpha helices and beta sheets, are consistent across the majority of *Sw*ASSTs, *Sw*ASST11 deviates slightly, displaying two fewer beta sheets at the N-terminus, likely due to its shorter length. Additionally, a beta sheet spanning residues 318-323 in *Sw*ASST1 is oriented in the reverse direction compared to its counterparts in other *Sw*ASST enzymes.

**Fig 12.**
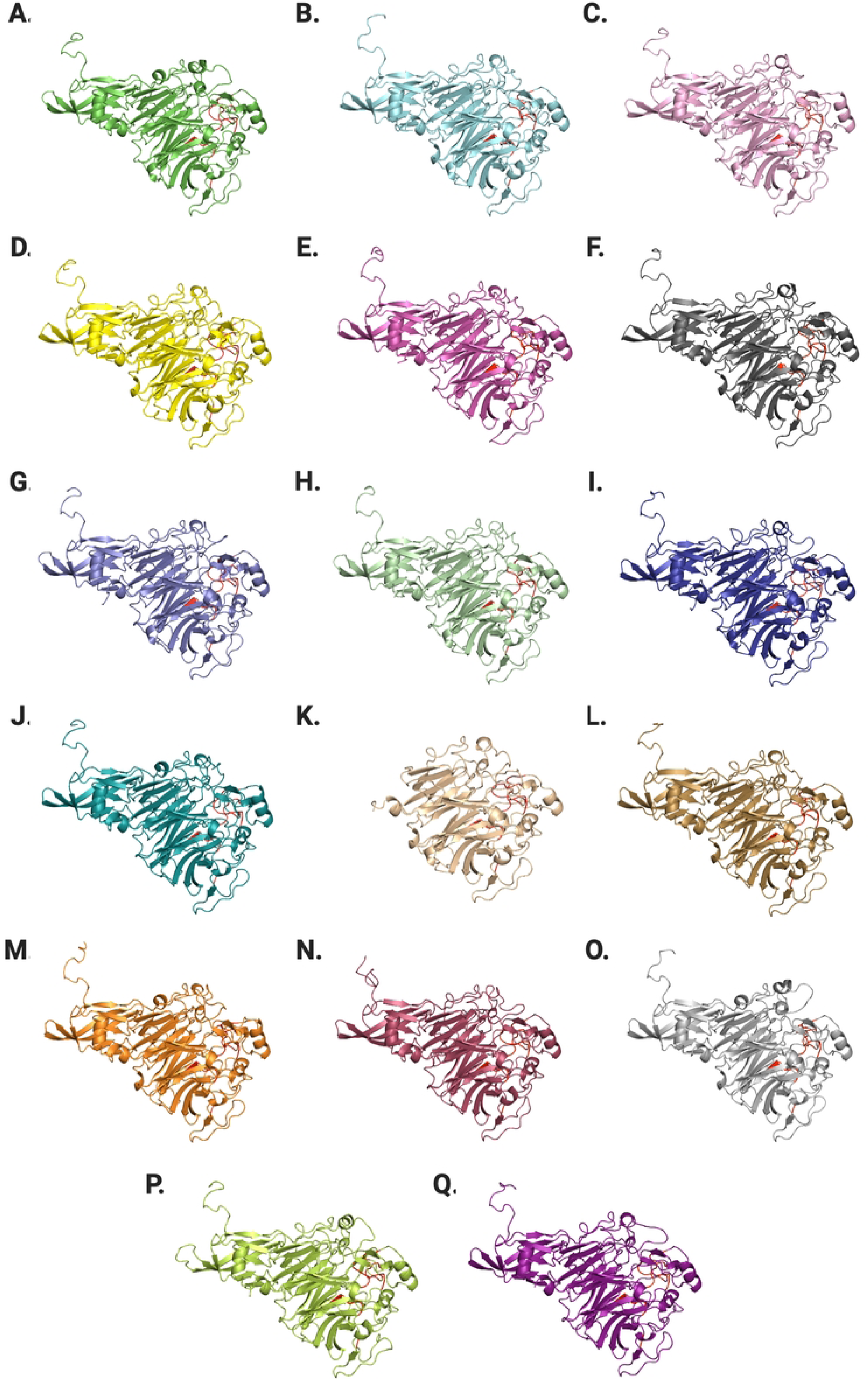
Structures of *Sw*ASST proteins generated by Alphafold2. From A to Q are predicted structures of *Sw*ASST1 to *Sw*ASST17. Conserved regions harboring active site residues are highlighted in red.

**Fig 13.**
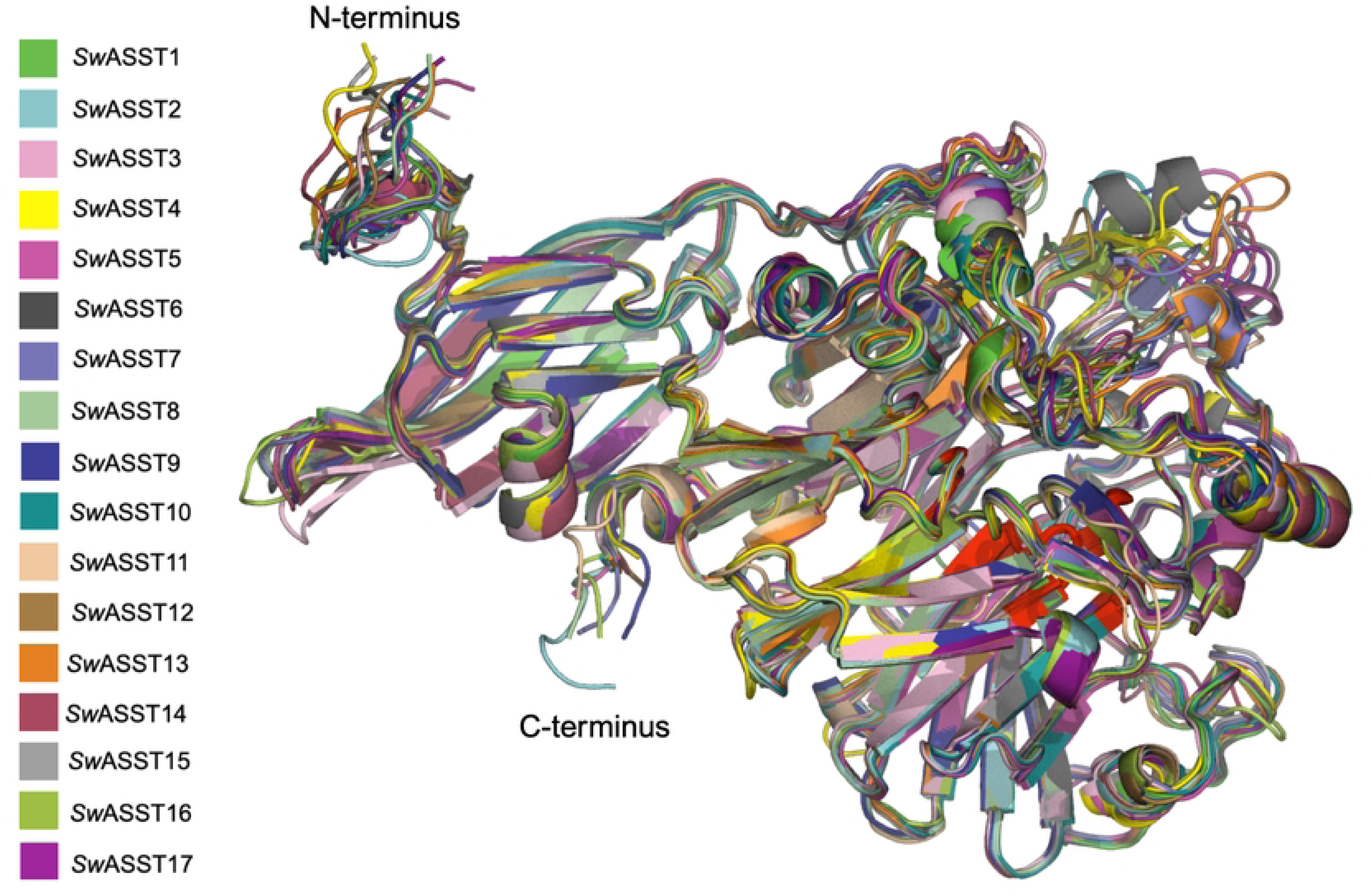
Structural alignment of *Sw*ASST proteins. The figure depicts aligned Alphafold2 derived structures of *Sw*ASSTs with conserved regions highlighted in red.

As can be seen in Fig 13, most areas of *Sw*ASSTs are completely superimposable and very well aligned. However, there are some regions where we see lower degree of structural alignment (Fig 14, blue and green areas). It is interesting to note that the areas with variable structural alignments can have either high or low amino acid sequence conservation. One such region is at the N-terminus with the conserved motif V/s/t-W-N-N-P-X-G-G-A-L/m/v-E-W (Fig 14, blue cluster, S3 Fig), displays considerable variation in the loop positions among the aligned *Sw*ASST structures. Contrastingly, at the C-terminus which is the other variable region (Fig 14, green cluster, S3 Fig), the level of structural alignment diminishes relative to the rest of the *Sw*ASST structures. This region is characterized by low sequence conservation and exhibits the most significant variability in secondary structural elements across the *Sw*ASST family.

**Fig 14.**
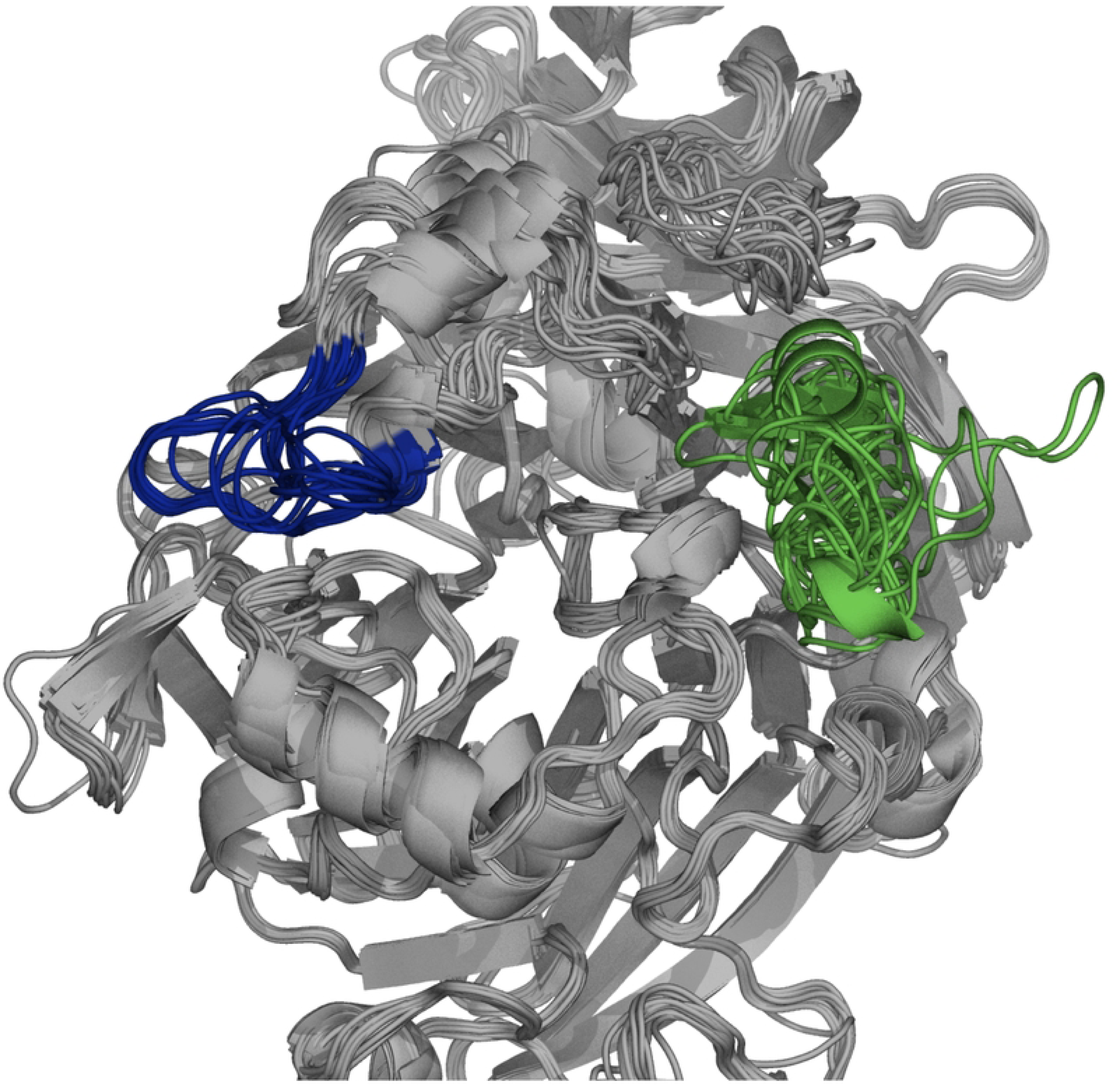
Unique clusters in *Sw*ASSTs alignments. Dark blue region represents the N-terminus cluster, and the C-terminus cluster is depicted in green. These regions show lower degree of structural alignments across the *Sw*ASSTs.

Sequence similarity studies and phylogenetic analyses indicate that *Sw*ASST3 is evolutionarily closest to *E. coli* ASST (*Ec*ASST), while *Sw*ASST6 is the most divergent (Fig 11B). Structural comparisons of *Ec*ASST(29) with *Sw*ASST3 and *Sw*ASST6 exhibit a substantial degree of alignment across the three structures (Fig 15). However, a loop region spanning residues 151-164 present in *Ec*ASST is absent in *Sw*ASST3 and *Sw*ASST6. Additionally, *Sw*ASST6 uniquely features an alpha helix and a loop between residues 555-568. We also included a ligand bound structure of *Ec*ASST in our alignment (Fig 15). In the ligand-bound structure of *Ec*ASST (green), the active site binds a molecule of para-nitrophenol (pNP), a reaction product of *Ec*ASST with the sulfate donor para-nitrophenyl sulfate (pNPS). Fig 16 presents a detailed view of the active site, showing the ligand bound within *Ec*ASST and the catalytic histidine residues from all three structures. While the histidine residues from the two *E. coli* structures align perfectly with each other, the histidine residues from the *Sw*ASSTs align amongst themselves but show slight positional differences when compared to the catalytic histidine residue of *Ec*ASST. This observation suggests subtle variations in the active site residue positioning in *Sw*ASSTs relative to *Ec*ASST.

**Fig 15.**
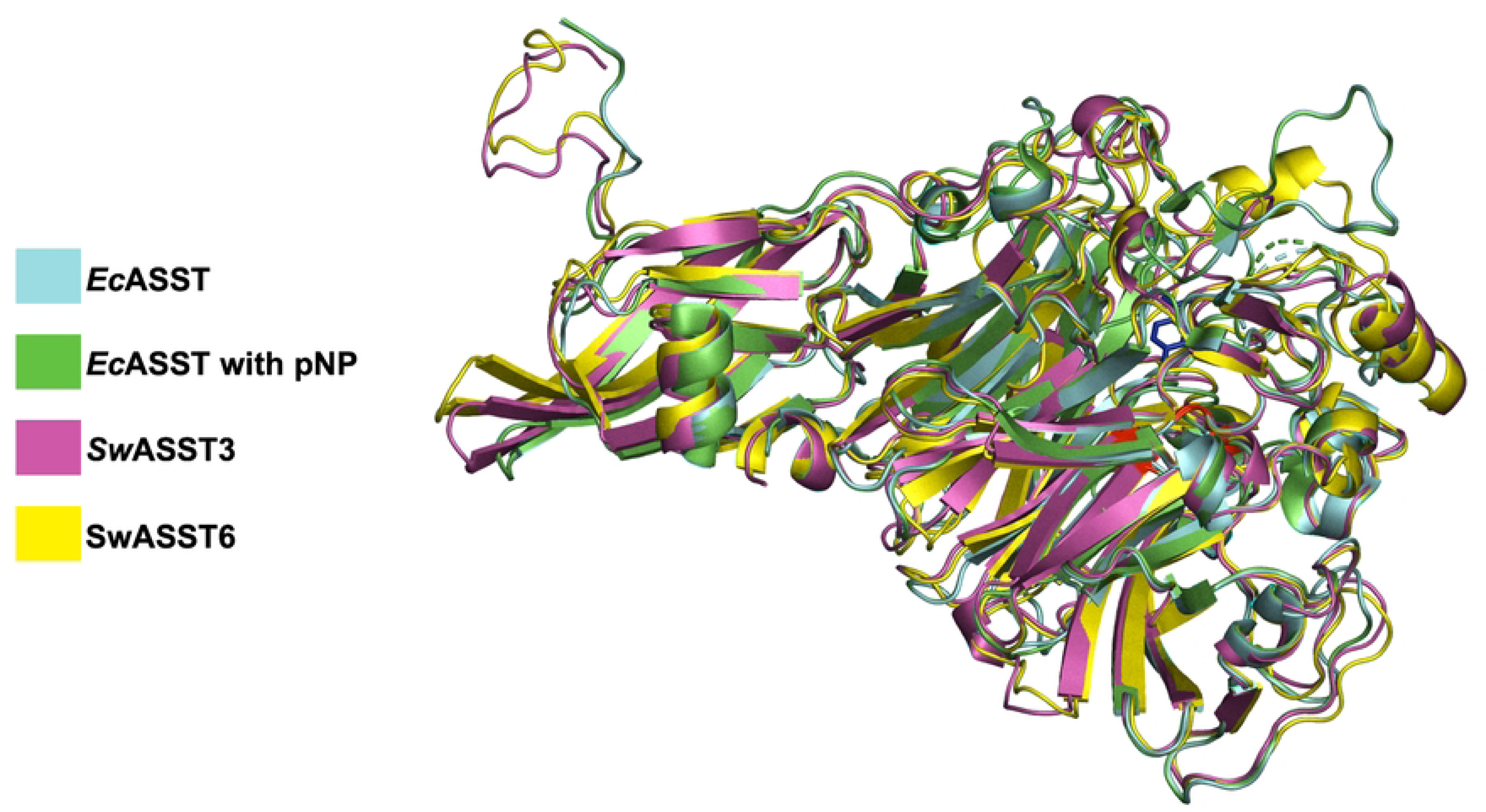
Alignment *Sw*ASST3 and *Sw*ASST6 with *Ec*ASST. The figure shows aligned structures of *Sw*ASST3 (pink) and *Sw*ASST6 (yellow) with free (cyan, PDB ID: 3ELQ) and ligand bound (green, PDB ID: 3ETT) structures of *Ec*ASST. There is a molecule of para-nitrophenol (shown as sticks in blue color) bound in the active site of *Ec*ASST (green, PDB ID: 3ETT).

**Fig 16.**
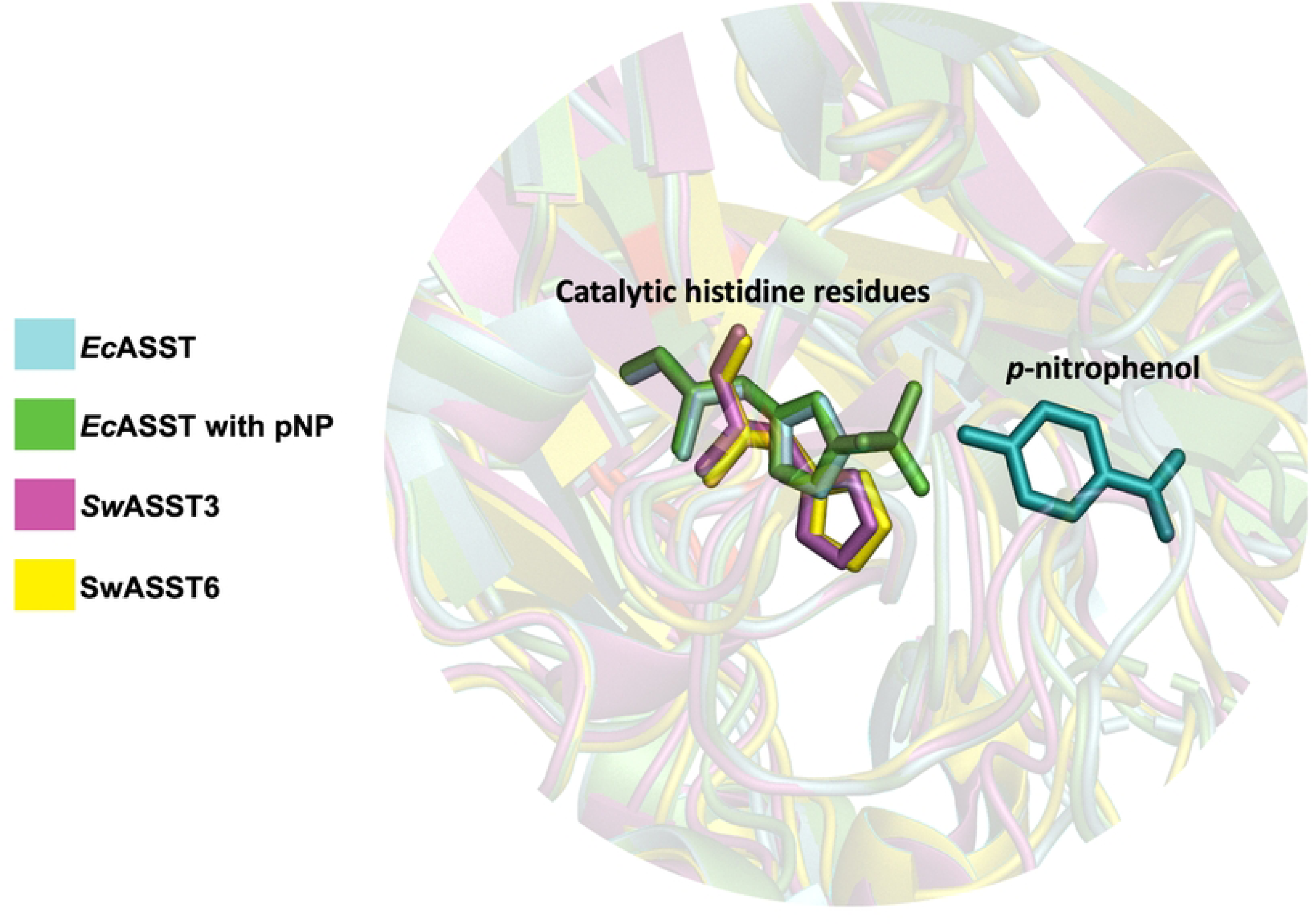
Active site of *Sw*ASSTs (3 and 6) and *Ec*ASST highlighting the catalytic histidine residues. The figure displays a zoomed in active site of *Sw*ASST3 (pink), *Sw*ASST6 (yellow), *Ec*ASST (cyan, free), and *Ec*ASST (green, pNP bound). Catalytic histidine residues are presented as sticks and highlighted in the same colors for each of the corresponding structures. pNP is shown as sticks (in cyan).

### Different classes of sulfotransferases from the members of the human gut microbiome and their comparison to human sulfotransferases

Sulfotransferases have historically been categorized based on their dependency on 3’-phosphoadenosine 5’-phosphosulfate (PAPS) as either PAPS-dependent, using PAPS as a sulfate donor, or PAPS-independent(29). Human sulfotransferases, typically around 300 amino acids in length, exclusively use PAPS, whereas microbial aryl-sulfate sulfotransferases (ASSTs), also known as PAPS-independent sulfotransferases, are generally larger. Recent findings, however, have identified PAPS-dependent sulfotransferases in gut commensals; these microbial enzymes are longer than their human counterparts, averaging around 370 amino acids, yet smaller than ASSTs(21, 22). To understand the prevalence of sulfotransferase classes, we explored the genomes of dominant human gut microbes, using protein blast searches with either *Sw*ASST1 (a PAPS-independent aryl-sulfate sulfotransferase) or *BT*_0416 (a PAPS-dependent cholesterol sulfotransferase from *Bacteroides thetaiotaomicron*) as references. The results demonstrate a wide distribution of both sulfotransferase types across gut microbiome members (S4 Fig). Specifically, the *Bacteroides* genus predominantly carries PAPS-dependent sulfotransferases, with *Parabacteroides* following closely (S5 Fig) whereas PAPS-independent sulfotransferases (aryl-sulfate sulfotransferases) are abundant in *Sutterella* (S6 Fig). Sequence (S7 Fig) and structural (S8 Fig) alignments were performed to elucidate the differences between these enzymes. The sequence similarity between *Sw*ASST1 and other PAPS-dependent sulfotransferases (*BT*_0416, hSULT1A1, and hSULT2B1) is very low. Within the PAPS-dependent class, however, there is higher sequence conservation. Structurally, there is a significant divergence between PAPS-independent (*Sw*ASST1) and PAPS-dependent sulfotransferases (S7 Fig, A). Human PAPS-dependent sulfotransferases align closely with each other and are completely superimposable (S7 Fig, B). When aligning all three PAPS-dependent sulfotransferases, including two from humans (hSULT1A1, and hSULT2B1) and one from gut microbes (*BT*_0416), some overlapping regions are apparent, but the structures are not completely superimposable (S7 Fig, C), particularly a large alpha-helix in *BT*_0416 that does not align with the others. A striking contrast is observed in the secondary structure composition between these two types of sulfotransferases. PAPS-dependent sulfotransferases are rich in alpha helices (S7 Fig, B and C), whereas PAPS-independent *Sw*ASSTs feature beta sheets as their dominant structural elements (Fig 12). This contrast suggests a divergent evolutionary path for these two classes of sulfotransferases.

## Conclusion

Sulfotransferases from *Sutterella wadsworthensis 3_1_45B* exhibit a mixture of shared and unique structural characteristics. While these enzymes generally align with a common structural fold, certain regions display variability in both sequence conservation and structural configuration. Sequence analysis reveals modest sequence similarity among *Sw*ASSTs, suggesting functional diversification despite a high degree of conservation in the active site. The presence of signal peptides in the majority of *Sw*ASSTs indicates potential translocation across the cytoplasmic membrane, with variations in sequence pointing to differential secretion pathways. Moreover, the analysis of transmembrane regions highlighted a subset of *Sw*ASSTs harboring predicted TM domains, which indicates a possible role in membrane integration or association. This observation, coupled with the presence of various secretion system components within the *S. wadsworthensis* genome, suggests that these sulfotransferases could be substrates for complex secretion pathways, potentially contributing to the metabolic functions within the human gut microbiome. Additionally, structural alignments and phylogenetic comparisons indicate both conservation and divergence from other well-characterized sulfotransferases, highlighting the complexity and evolutionary plasticity of these enzymes.

## Methods

### Members of the human gut microbiome harboring annotated *asst* genes

IMG Genome was used to search for the prevalent human gut microbes(90, 91) that contain annotated genes for *asst* that are predicted to produce the protein aryl-sulfate sulfotransferases (ASSTs). Through a search of IMG genome database with either enzyme name, aryl-sulfate sulfotransferases or the enzyme ID, EC 2.8.2.22, produced lists of genomes containing predicted *asst* genes. By selecting genome name under the filter column, lists of predicted ASSTs for each specific genus were collected. The total number of sulfotransferase genes present in a genus were counted by adding annotated genes from all the species under that genus.

### Annotated *asst* genes from the genus *Sutterella*

By selecting genus *Sutterella* in the IMG genome database, we were able to collect different species and strains harboring annotated *asst* genes. From this, *Sutterella wadsworthensis 3_1_45B* was selected due to the number of annotated sulfotransferases found in its genome. Locus tags, gene IDs, GC content, and gene neighborhoods were retrieved for all annotated *asst* genes via IMG genome database.

### Properties of predicted ASST proteins from *Sutterella wadsworthensis 3_1_45B*

Amino acid sequences for all 17 predicted ASST proteins from *S. wadsworthensis 3_1_45B* were obtained from the IMG genome database and UniProt and were aligned to confirm that amino acid sequences retrieved from both sources are exactly same.

#### Signal peptides and transmembrane regions

Amino acid sequences for all predicted sulfotransferases were analyzed with SignalP 6.0 and TMHMM 2.0 using preset parameters to gain insights into the presence and variability of signal peptides and transmembrane helices respectively. These collected datasets were also verified with IMG genome database. In addition to providing information about presence or absence of signal peptides, SingalP 6.0 also allowed for identification of the possible signal peptide type, along with the length of the signal.

#### Multiple Sequence Alignment

Multiple sequence alignment for all 17 ASST amino acid sequences was performed with the help of UniProt alignment tool, which utilizes the Clustal Omega program 2.1. This analysis provided information about sequence conservation in predicted ASST proteins. Additionally, percent identity matrix for these 17 ASST sequences was obtained from the alignments. Multiple alignments were performed with and without the signal peptides. Alignments with signal peptides were generated to understand differences in these regions. Alignments without signal peptides were generated to understand the variability in ASST sequences.

#### ASST structure predictions with AlphaFold2

Amino acid sequences of ASSTs without signal peptides were uploaded to Colaboratory AlphaFold2 (ColabFold v1.5.3: AlphaFold2 using MMseqs2) and default settings were used for all runs to predict protein structures(92, 93). The three conserved regions found from the multiple sequence alignment of *Sw*ASSTs with *Ec*ASST were highlighted in red in all ASST structures.

#### ASST structural alignments with PyMOL

The PyMOL alignment tool with the default parameters and align command with five iteration cycles and a cutoff of 2 Å was utilized to align all the structures except for the alignment ‘c’ described below. These alignments are: a) For the alignment among all 17 ASSTs. All structures were aligned to *Sw*ASST1. b) Alignment of *E. coli* ASST (*Ec*ASST) with *Sw*ASST3 and *Sw*ASST6 where two separate PDB structures of *Ec*ASST were utilized. Structure with PDB ID 3ETT has para-nitrophenol (pNP) bound to the active site, while 3ELQ has no bound ligands in the active site. c) Alignment of *Sw*ASST1 with *Bacteroides thetaiotaomicron* VPI-5482 sulfotransferase *BT*_0416 (structure generated via AlphaFold2), human sulfotransferase 1A1 (hSULT1A1, PDB ID 1LS6), and human sulfotransferase 2B1b (hSULT2B1b, PDB ID 1Q1Z). hSULT2B1b, hSULT1A1, *BT*_0416, and *Sw*ASST1 were aligned using the cealign command in PyMOL’s alignment tool due to low sequence similarity.

### Search for PAPS-dependent and PAPS-independent sulfotransferases in the members of the human gut microbiome

Using NCBI protein blast, two separate blasts, one with a sulfotransferase from *Bacteroides thetaiotaomicron* sulfotransferase, *BT*_0416 and another one with an aryl sulfate sulfotransferase from *Sutterella wadsworthensis 3_1_45B*, *Sw*ASST1 were performed against members of the human gut microbiome. *BT*_0416 is PAPS-dependent sulfotransferase and *Sw*ASST1 is a PAPS-independent sulfotransferase. *Bacteroides* and *Sutterella* were each excluded from their respective blast searches. Each genus of gut microbes was entered into the filter bar to determine if that genus contained PAPS-dependent and/or PAPS-independent sulfotransferases. From this search a binary dataset was created where presence was marked by 1 and absence was marked by 0. This dataset was utilized to create a heatmap. Predicted sulfotransferase proteins from different species and strains of each genus were added together to calculate the total number of predicted sulfotransferases (PAPS-dependent or independent) per genus.

## Supporting information

**S1 Fig. Partial multiple sequence alignment of all 17 *Sw*ASSTs with *Ec*ASST.** Sequence alignment was performed on protein sequences without signal peptides. Color bar above aligned sequences shows the amino acid conservation at that position where dark red indicates highly conserved residues, while dark blue indicates little to no conservation.

**S2 Fig. IDDT plots for all predicted structures of *Sw*ASST proteins generated by Alphafold2.** Each plot has confidence percentage on the y-axis and the amino acid position on the X-axis.

**S3 Fig. N and C terminus regions showing variable structural alignment.** N and C terminus regions of *Sw*ASSTs that show high variations during structural alignments. A) Sequence alignment highlighting the N-terminus region that shows sequence conservation but structural variation. B) C-terminus multiple sequence alignment highlighting region with the red box that shows high secondary structure variability.

**S4 Fig. Distribution of PAPS-dependent and PAPS-independent sulfotransferases across members of the human gut microbiome.** Heat map representation for the presence of sulfotransferase that are similar to *Sw*ASST1 (PAPS-independent) and/or *BT*_0416 (PAPS-dependent). Green boxes indicate the presence of sulfotransferases similar to *Sw*ASST1 whereas blue boxes indicate sulfotransferases similar to *BT*_0416.

**S5 Fig. Human gut microbial genera with sulfotransferases similar to *BT*_0416.** The bar graph represents total number of annotated sulfotransferases similar to *BT*_0416 in species and strains from each genus.

**S6 Fig. Human gut microbial genera with sulfotransferases similar to *Sw*ASST1.** The bar graph represents total number of annotated sulfotransferases similar to *Sw*ASST1 in species and strains from each genus.

**S7 Fig. Multiple sequence alignment of *Sw*ASST1 with *Bacteroides BT*_0416, hSULT1A1, and hSULT2B1b.** Light blue highlighted regions are well conserved regions and regions that share a high percentage of similarity. Color bar above the sequences show the amino acid conservation at that position, where dark red is the most conserved and dark blue is the least conserved position.

**S8 Fig. Structural alignments of *Sw*ASST1 with *BT*_0416, hSULT1A1, and hSULT2B1b.** A) Alignment of all four sulfotransferases. B) Alignment of both human sulfotransferases, hSULT1A1 and hSULT2B1b. C) Alignment of three PAPS-dependent sulfotransferases (*BT*_0416, hSULT1A1, and hSULT2B1b). PAPS is shown in red and pNP is in lime green (visible in alignment B).

**S1 Table**. **Human sulfotransferases and their known substrates (sulfate acceptors).** Shown are the lengths of each human sulfotransferase enzyme along with its known substrate, and UniProt ID number.

**S2 Table**. **Annotated *asst* genes in the genus *Sutterella***. Table depicts the information shown in Fig 3 which includes total number of annotated asst *genes* from different species and strains of *Sutterella*.

**S3 Table**. **Aryl-sulfate sulfotransferase enzymes from *S. wadsworthensis 3_1_45B.*** The table shows properties of each annotated *asst* gene and its protein product.

**S4 Table. Analysis of transmembrane helices from *Sw*ASSTs.** Lists architectural information of the predicted TM regions.

